# Soma and Neurite Density MRI (SANDI) of the in-vivo mouse brain

**DOI:** 10.1101/2021.08.11.455923

**Authors:** Andrada Ianuş, Joana Carvalho, Francisca F. Fernandes, Renata Cruz, Cristina Chavarrias, Marco Palombo, Noam Shemesh

**Author notes:** Corresponding author: Dr. Noam Shemesh, Champalimaud Research, Champalimaud Centre for the Unknown, Av. Brasilia 1400-038, Lisbon, Portugal.; ext. #4467. Dr. Andrada Ianus, Champalimaud Research, Champalimaud Centre for the Unknown, Av. Brasilia 1400-038, Lisbon, Portugal.

## Abstract

Diffusion MRI (dMRI) provides unique insights into the neural tissue milieu by probing interaction of diffusing molecules and tissue microstructure. Most dMRI techniques focus on white matter tissues (WM) due to the relatively simpler modelling of diffusion in the more organized tracts; however, interest is growing in gray matter characterisations. The Soma and Neurite Density MRI (SANDI) methodology harnesses a model incorporating water diffusion in spherical objects (assumed to be associated with cell bodies) and in impermeable “sticks” (representing neurites), which potentially enables the characterisation of cellular and neurite densities. Recognising the importance of rodents in animal models of development, aging, plasticity, and disease, we here sought to develop SANDI for preclinical imaging and provide a validation of the methodology by comparing its metrics with the Allen mouse brain atlas. SANDI was implemented on a 9.4T scanner equipped with a cryogenic coil, and experiments were carried out on N=6 mice. Pixelwise, ROI-based, and atlas comparisons were performed, and results were also compared to more standard Diffusion Kurtosis MRI (DKI) metrics. We further investigated effects of different pre-processing pipelines, specifically the comparison of magnitude and real-valued data, as well as different acceleration factors. Our findings reveal excellent reproducibility of the SANDI parameters, including the sphere and stick fraction as well as sphere size. More strikingly, we find a very good rank correlation between SANDI-driven soma fraction and Allen brain atlas contrast (which represents the cellular density in the mouse brain). Although some DKI parameters (FA, MD) correlated with some SANDI parameters in some ROIs, they did not correlate nearly as well as SANDI parameters with the Allen atlas, suggesting a much more specific nature of the SANDI parameters. We conclude that SANDI is a viable preclinical MRI technique that can greatly contribute to research on brain tissue microstructure.

## 1. Introduction

The development of non-invasive imaging biomarkers which can reflect microscopic tissue properties, for instance the size and density of cells and/or neurite projections [1, 2], is a topic of great interest for neuroimaging in general and diffusion Magnetic Resonance Imaging (dMRI) in particular. Extracting such microstructural information can be valuable for assessing changes upon brain development [3], plasticity [4], or aging [5], as well as for the diagnosis and monitoring of various neurologic, neurodegenerative, and psychiatric diseases [5, 6], as well as neural injury such as stroke [7, 8] [9] or traumatic brain injury[10], cancer infiltration [11], and other conditions.

Diffusion Magnetic Resonance Imaging (dMRI) has become one of the most valuable modalities for probing tissues noninvasively, with sensitivity to the microscopic scale. In dMRI, the signal is sensitized to the displacement of water molecules in the tissue, which is strongly influenced by microscopic boundaries imparted by cellular and subcellular structures [12]. Various approaches have been proposed in the literature to characterise dMRI signals [1, 2, 13, 14] and can be considered as part of two broad categories. *Signal representations*, on one hand, employ general functional forms of the dMRI signal decay, for example by considering different orders of the cumulant expansion, with diffusion tensor imaging (DTI) [15] and diffusion kurtosis imaging (DKI)[16] being prominent examples. While precise, these methods are not considered very specific since the underlying contributions to the metrics are not resolved, and since many different tissue features can lead to similar signal decays [17, 18]. On the other hand, to characterize tissue features more specifically, *biophysical modelling* approaches aim to disentangle the signal contributions from two or more water pools, usually assigned to different tissue components, such as intra- and extra-cellular space [1] [2, 13, 19–21]. When thoroughly validated, these models can be considered more specific than representations (and often require more measurements to resolve the microstructural features) [22–30].

Most brain tissue models employed with dMRI data make the explicit assumption of exactly two compartments, e.g. [22–24, 31, 32], with the following diffusion properties: signal ascribed to the intra-neurite space (i.e. axons and dendrites) is assumed to be described by diffusion inside impermeable cylinders, usually considered to have zero radius (i.e. “sticks” [31]), while the extra cellular space is assumed to be described by signal exhibiting either isotropic or anisotropic Gaussian diffusion [13, 21–24, 31–34]. Thus, the Gaussian diffusion component models signal from water in the extra-cellular space as well inside cell bodies. At high and very high b-values, such a model would yield a well characterized power-law of the directionally averaged data as the b-value is increased. However, recent studies show a departure from this behaviour at (very) high b-values, which has been used to inform modelling approaches, for instance to characterise axon diameter in white matter [35], or to include an additional compartment of slow diffusion [36, 37]. In some studies, the slow diffusion compartment is represented by diffusion restricted in spheres [37] and has been proposed to model the signal from cell bodies (and other quasi spherical structures), with a significant contribution especially in gray matter (GM) [38–41].

The recently-introduced Soma and Neurite Density Imaging (SANDI) methodology [39] aims to characterise such spherical object contributions using standard single diffusion encoding (SDE) dMRI, acquired with several shells (usually ≥ 5, depending on the number of fitted parameters[42]) up to (very) high b-values and powder-averaged data. The SANDI model relies on a three-compartment model consisting of sticks, spheres and isotropic Gaussian diffusion fitted to the powdered averaged data. Insofar, SANDI has been applied to in-vivo human brain imaging on a high-performance scanner, as well as ex-vivo in the mouse brain, where a preliminary comparison with histology data showed a good correlation between the soma signal fraction and signal intensity of DAPI stain, a marker for cell nuclei [30]. Nevertheless, estimating the spherically restricted signal fraction is challenging due to signal-to-noise and contrast-to-noise limitations as well as bias due to Rician noise floor [40, 43]. Other studies have combined standard SDE acquisitions with more general B-tensor encoding schemes showing a potential benefit of multi-waveform approaches [40, 41], nevertheless, such sequences are not widely available, neither on clinical nor on pre-clinical systems.

In this study we aim to address several gaps in the evaluation and application of the original SANDI approach based on SDE acquisitions: i) we investigate the stability of SANDI metrics in the mouse brain, in-vivo at 9.4T; ii) we utilise the complex dMRI data to assess experimentally the effects of Rician noise floor on the SANDI parameters; iii) we compare parameters derived from SANDI and a more established technique, namely Diffusion Kurtosis Imaging (DKI), and iv) assess their correspondence to a histological proxy of cell density based on the Allen Brain atlas; v) lasty, we also investigate the possibility of reducing the number of shells in the diffusion protocol and its impact on the estimated parameters.

## 2. Methods

### 2.1 In-vivo MRI experiments

All animal studies were approved by the competent institutional and national authorities, and performed according to European Directive 2010/63.

In-vivo dMRI data was acquired from N=6 C57BL/6J mice (N=1 males, N=5 females, 34 ± 5 weeks old, grown in a 12 h/12 h light/dark cycle with ad libitum access to food and water) on a 9.4T Bruker Biospec scanner equipped with an 86 mm quadrature transmission coil and 4-element array reception cryocoil. Briefly, mice were induced with 5% isoflurane mixed with oxygen-enriched medical air and maintained at 1.5-2%, and their temperature and breathing rate were continuously monitored.

Diffusion MRI datasets for SANDI were acquired using a PGSE-EPI sequence with the following parameters: TE = 36.8 ms, TR = 4 s, 4 averages, slice thickness = 0.4 mm, 35 slices, in plane resolution = 0.12 × 0.12 mm^2^, FOV = 14.2 × 12 mm^2^, matrix = 118 × 100, Partial Fourier = 1.35, per-slice triggering and with fat suppression. The EPI acquisition bandwidth was 375 kHz and data were acquired in a single shot.

In terms of diffusion weighting, the data has 8 shells with b-values of 1, 2.5, 4, 5.5, 7, 8.5, 10 and 12.5 ms/μm^2^ and 40 directions each, distributed on a hemisphere following the default directions from the manufacturer. The diffusion time Δ = 20 ms was chosen to provide sensitivity for mapping apparent soma density according to previous simulation studies [43], and the gradient duration δ = 5.5 ms was chosen based on the maximum diffusion weighting given the hardware constraints (max gradient strength of ^~^ 660 mT/m). The acquisition also included anatomical images acquired with a RARE sequence: RARE factor 8, TE = 36.2 ms, TR = 3.2 s, 4 averages, slice thickness = 0.4mm, 35 slices, in plane resolution = 0.08 × 0.08 mm, matrix = 178 × 150, Partial Fourier = 1.1, per-slice triggering and fat suppression. The acquisition time for this protocol including all necessary adjustments was ^~^ 2.5h.

### 2.2 Data pre-processing

Complex data for each of the four receiver channels was exported from the scanner and processed in Matlab^®^ (The MathWorks, Natick, MA, USA), unless otherwise specified, and the pre-processing pipeline is illustrated in Figure 1.

**Figure 1.**
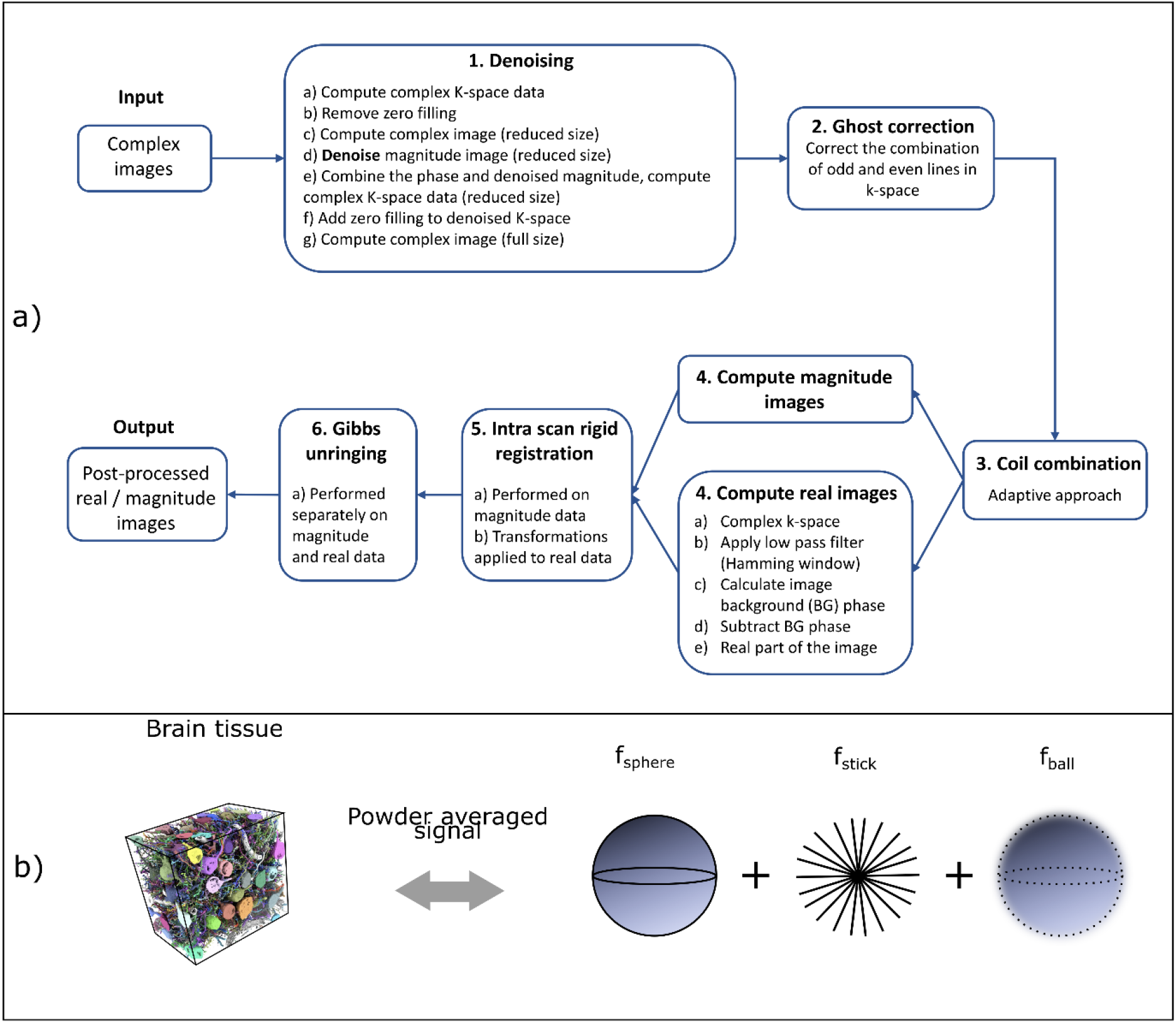
a) Schematic description of the data processing pipeline. Input: complex images for each individual channel after the Bruker reconstruction from the scanner. Step 1: images are denoised for each channel. Step 2: images are ghost corrected by accounting for phase differences between odd and even k-space lines. Step 3: the four channels are combined following an adaptive approach. Step 4: magnitude and real images are computed. Step 5: images are registered within each scan. Step 6: images are corrected for Gibbs ringing. Output: Post-processed real and magnitude images. b) Example of mouse brain tissue reconstructed based on electron microscopy reproduced with permission from [53] (left) and schematic representation of the SANDI model (right). The powder averaged diffusion signal is a combination of diffusion restricted in spheres with a signal fraction f_stick_, diffusion restricted in sticks with signal fraction f_sticks_ and isotropic Gaussian diffusion with signal fraction f_ball_, with f_sphere_ + f_stick_ + f_ball_ = 1.

#### Step 1

The first step was denoising the complex data using the Marchenko-Pastur PCA (MP-PCA) denoising technique described in [44] and implemented in Matlab^®^[45]. As noise correlations in the image, due to data combination from multiple channels as well as zero filling in k-space (which is automatically performed by the vendor as part of the reconstruction pipeline), would negatively impact the performance of the denoising, we performed the following steps on the data: a) for the complex image for each channel we calculated the complex k-space data using a 2-dimensional inverse Fourier Transform, b) then we removed the zero padding and c) computed the (slightly lower resolution) complex image. Then, d) we denoised the magnitude of the lower resolution image, as the application of the MP-PCA denoising to magnitude data is well characterized, e) then we combined it with the phase information and calculated the corresponding denoised complex k-space. f) Next, we substituted the measured k-space data with the denoised one, and g) finally, we computed the denoised complex image at the original resolution.

#### Step 2

The next step was ghost correction based on the algorithm described in [46], to correct the phases differences between odd and even k-space lines from EPI acquisition. We used an inhouse Matlab^®^ implementation of the algorithm.

#### Step 3

Following the ghost correction, the complex data from the 4 different channels was combined using the adaptive approach described in [47, 48] implemented in Matlab^®^[49].

#### Step 4

At this step we calculated the standard magnitude images by taking the absolute values of the complex data, as well as the real images after the background phase removal. To obtain the real data, we followed the steps described in [50, 51]. a) First, we calculated the complex k-space, then b) we a applied a low pass filter, specifically, a Hamming window centred on the k-space centre and with a width equal to the amount of symmetrically acquired k-space lines, as described in [50]. c) Next, the background phase after applying the low pass filter was calculated and d) subtracted from the image phase; e) the resulting real part is used in subsequent analyses and referred to as “real image”.

#### Step 5

The last step of this processing pipeline was the intra-scan registration (3D rigid registration based on mutual information), necessary due to the long duration of the scans to account for any movement as well as effects from gradient heating. As the diffusion data was measured in two acquisitions (8 b = 0 ms/μm^2^ volumes followed by odd and even b-values, respectively), with the first loop over b-values and the second over directions (i.e. b1 dir1, b2 dir1, …, b1 dir2, b2 dir2, …), the registration was performed as follows: a) first, the measurements with the lowest b-values (i.e. b = 1 ms/μm^2^ for the first acquisition and b = 2.5 ms/μm^2^ for the second one) were registered to the first direction, then the transformation was applied to the higher b-values from the respective direction; b) the b = 0 ms/μm^2^ images were registered to the last one, then c) this reference b = 0 ms/μm^2^ image was registered between the second and first acquisition sets, and d) this transformation was applied to all diffusion weighted volumes. The registration was performed using the *imregister* function in MATLAB on magnitude images and the same transformations were applied to real data.

#### Step 6

After registration, real and magnitude data were individually corrected for Gibbs ringing, using a Matlab^®^ implementation of the algorithm proposed by Kellner et al [52].

Finally, the data was normalized by the mean of the b = 0 ms/μm^2^ images, and for the SANDI analysis, the diffusion weighted images were averaged over directions for each shell to calculate the powder averaged signal.

### 2.2 SANDI analysis

In the first analysis, we employ the SANDI framework to analyse the powder-averaged diffusion data, as detailed in [39].

#### 2.2.1 Model formulation

The SANDI model assumes the diffusion signal can be explained by three compartments, namely intra-neurite signal which is modelled as diffusion inside impermeable sticks with zero radial diffusivity, intra-soma signal which is modelled as restricted diffusion inside spheres, and extra-cellular signal, which is modelled as Gaussian diffusion, as illustrated in Figure 2. The powder-averaged normalized diffusion signal has thus the following expression:

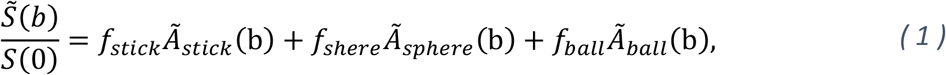

where 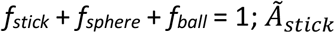 and 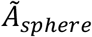 are the normalised, directionally averaged signals for restricted diffusion within neurites and soma, respectively and 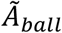 the normalised, directionally averaged signal of the extra-cellular space. The specific expressions are given bellow:

**Figure 2.**
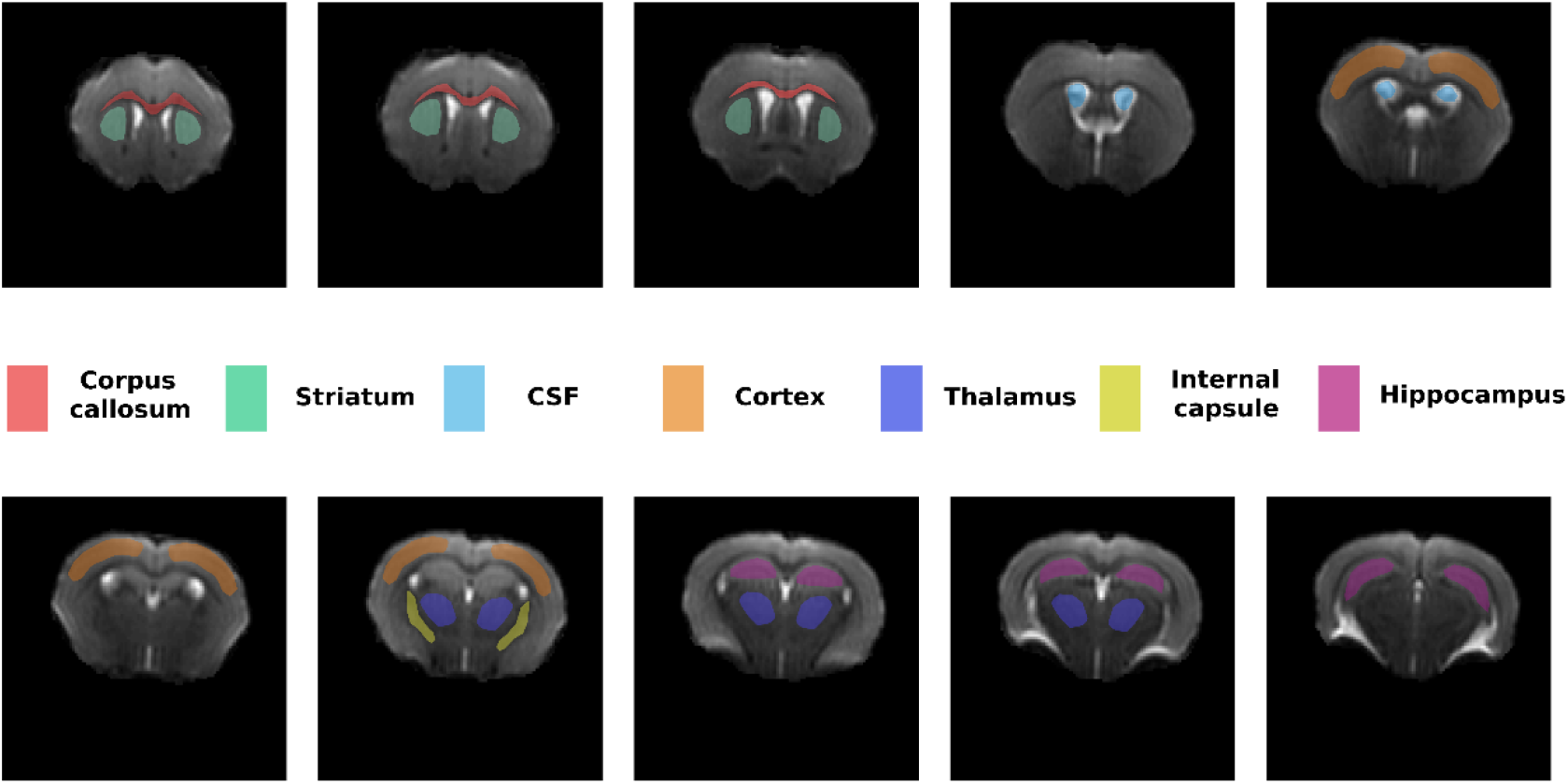
Graphical depiction of the different ROIs analysed.

##### Extra-cellular compartment

The diffusion of water molecules associated with the extra-cellular compartment, 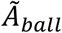 is modelled as isotropic Gaussian diffusion with a scalar effective diffusion constant *D_ball_*:

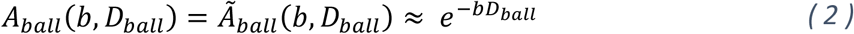

##### Intra-neurite compartment

The signal contribution 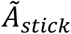 from neurites (dendrites and axons) assumes that neuronal processes can be described as a collection of long thin cylinders, with a parallel apparent diffusion coefficient 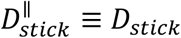 and a negligible perpendicular one 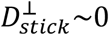. Under these assumptions, the direction-averaged 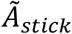 can be computed as powder average of randomly oriented sticks, such that [21, 54–59]:

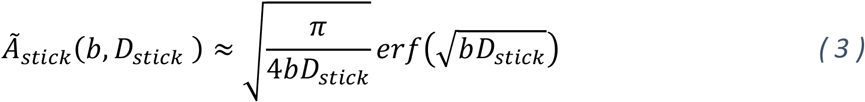

##### Intra-soma compartment

The signal contribution 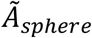 from cell bodies and other quasi spherical tissue components (astrocytes, glial cells, etc) is assumed to arise from a pool of diffusing water molecules restricted inside impermeable spheres with an effective radius *R_sphere_* and characterised by a diffusion coefficient *D_sphere_*. The normalized signal 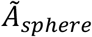 can be computed following the Gaussian Phase Distribution (GPD) approximation [60, 61], such that:

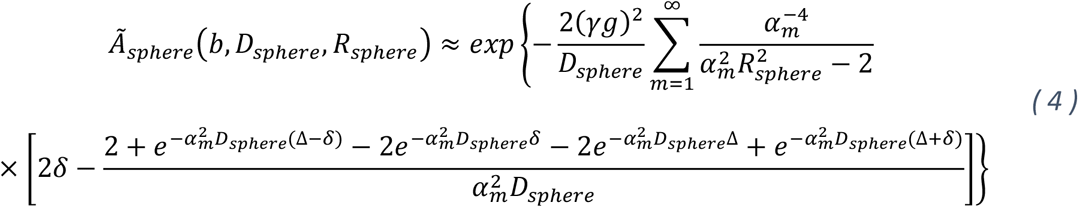

where *D_sphere_* is the bulk diffusivity of water inside the spherical compartment, δ and Δ the diffusion gradient pulse width and separation, g the magnitude of diffusion gradient pulse, α_m_ the m^th^ root of the equation 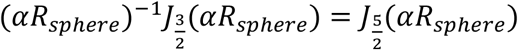, with J_n_(x) the Bessel function of the first kind. For simplicity, here we consider a single apparent radius *R_sphere_* as representative for all the somas, as a proxy for the volume weighted size distribution from a given MRI voxel, as detailed in [39]. For the diffusion weighting values and sphere radii considered here, the GPD provides a good proxy of the dMRI signal inside spheres [43, 62].

This model formulation assumes that on the timescale of the diffusion experiments (t_d_ ^~^ 20 ms), the effects of exchange between different compartments can be neglected. Moreover, it does not explicitly account for size distributions as well as for intra-compartment kurtosis, which is neglected in the GPD approximation.

#### 2.2.2 Model parametrization

Following equations (1)–(4), the parameters to be estimated from the direction-averaged data are: *D_stick_*, *D_sphere_*, *D_ball_*, *R_sphere_* as well as the signal fractions subject to the constraint *f_stick_* + *f_sphere_* + *f_ball_* = 1. For diffusion restricted in spheres, i.e. equation (4), the signal depends on the ratio *D_sphere_/R_sphere_*^2^, thus disentangling both the bulk diffusivity and restriction size from measurements which vary only the gradient strength is challenging. Thus, following the approach in [39], we fix the bulk diffusivity *D_sphere_* to 2 μm^2^/ms, a value measured for intracellular diffusivity at ultra-short diffusion times [63], and then we estimate *R_sphere_*. Moreover, it has been shown that fixing the diffusivities to slightly different values will not have a large impact on the pattern of estimated radii [64]. To ensure the signal fractions of the three compartments add up to 1, we have parametrized them as follows: *f_stick_* = *f_1_*, *f_ball_* = *f_2_*(*1-f_1_*) and *f_sphere_* = (*1-f_2_*)(*1-f_1_*), where f_1_ and f_2_ are between 0 and 1. Furthermore, to ensure f_1_ and f_2_ are indeed constrained between 0 and 1, they have been parametrized as cos(**α**_1_)^2^ and cos(**α**_2_)^2^, respectively, as proposed in [21].

#### 2.2.3 Radom Forest Regression

To estimate the five model parameters of SANDI we have employed a Random Forest (RF) regression algorithm [39]. Specifically, we have used the *TreeBagger* Matlab^®^ algorithm with 200 trees, i.e. the same number of trees as performed in the original SANDI analysis, and default settings for regression to estimate separately each SANDI parameter. Thus, a separate RF was trained for each parameter. The data for training consisted of the normalized and directionally averaged diffusion signal simulated using the same acquisition as the experimental data and 10^5^ SANDI parameter combinations, uniformly sampled as follows: *f_1_* and *f_2_ ∈[0, 1], D_stick_ ∈[0.25, 3]* μm^2^/ms, *D_ball_ ∈[0.25, 3]* μm^2^/ms and *R_sphere_ ∈[1, 10]* μm. To each datapoint, Gaussian noise with a standard deviation σ = 6.7E-3 was added, corresponding to an SNR of the powder averaged non-diffusion weighted data of 150. The value of σ was approximated based on the SNR in white matter as follows: the SNR of b_0_ images was ^~^25, with corresponding noise standard deviations of the normalized signal σ_0_ ^~^ 0.04. When averaging N images, the noise standard deviation of the averaged data decreases with 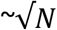, thus for the powder averaged data over 40 directions, the standard deviation of the noise was less than 6.7E-3, the value used in the training dataset.

### 2.3 DKI analysis

To further compare our results with a more established model of diffusion, we have also employed a diffusion kurtosis (DKI) analysis [16]. Thus, diffusion and kurtosis tensors were fitted voxelwise to the first 2 shells (b-values of 1 and 2.5 ms/μm^2^), that provide appropriate diffusion weighting for DKI [65], using a linear least squares algorithm implemented in MATLAB, employing the following equation:

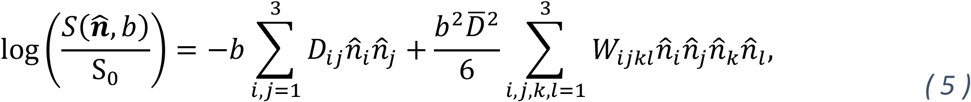

with *D_ij_* is the *ij* element of the symmetric rank 2 diffusion tensor **D** with trace 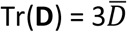, and *W_ijkl_* is the *ijkl* element of the symmetric rank 4 kurtosis tensor **W** and 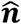 denotes the direction of the diffusion gradient. Next, scalar metrics, specifically mean diffusivity (MD), fractional anisotropy (FA) and mean kurtosis (MK), were calculated and included in the subsequent analyses, as follows [66]:

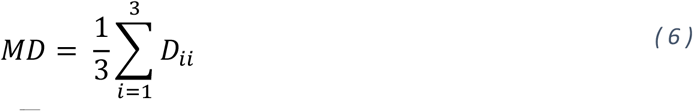

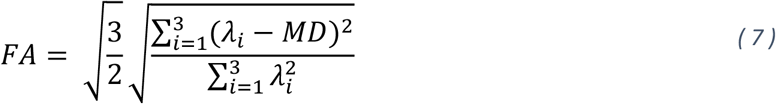

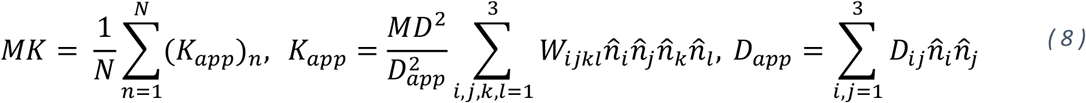

where *λ_i_* are the eigenvalues of the diffusion tensor, N is the total number of gradient directions and *D_app_* and *K_app_* are the apparent diffusion and kurtosis values along the n^th^ gradient direction.

### 2.4 Magnitude vs Real data

Magnitude data is in most cases the standard output from a dMRI acquisition and is used in many dMRI techniques. Nevertheless, the Rician distribution of the noise in the magnitude data can lead to parameter biases especially when including measurements with very high diffusion weighting close to the noise floor [40, 50], as performed in the SANDI technique. One way to remove this bias is to model the Rician noise floor and remove it [57] or, when complex data is available, employ real data instead [50, 51]. Thus, in the first analysis, we investigate the correlations of the SANDI parameters when obtained either from magnitude data, which is the standard approach, or real data, which provides less bias when averaging the signal [51], following the pre-processing pipeline described in Section 2.2. Given that the SANDI analysis uses the powder averaged data, to minimize the effect of the Rician noise floor on the estimated parameters[50, 51], in the subsequent analyses we use the parameters derived from the real data.

### 2.5 ROI analysis

To analyse the range of SANDI parameters in the mouse brain, in-vivo, we performed a region of interest (ROI) analysis including gray matter (GM) areas (cortex, thalamus, striatum, hippocampus), white matter (WM) areas (corpus callosum, internal capsule) and cerebrospinal fluid (CSF) regions. For each animal, the representative ROIs for each structure were manually delineated on the S_0_ images upon comparison with an atlas, and are illustrated in Figure 2.

The ROI analysis was employed to:

i. analyse the differences of SANDI parameters derived from magnitude and real data to investigate the effect of Rician noise in different parts of the brain. Correlation analysis was used to assess the agreement between the two cases, and Bland-Altman plots [67] were employed to better observe biases of the estimated parameters;
ii. investigate the parameter distributions across different areas of the brain. The parameter variability across animals was assessed by calculating the coefficient of variation of the mean ROI values across animals, both for SANDI and DKI parameters;
iii. analyse the correlations between SANDI and DKI parameters in different areas of the brain. From this analysis the CSF ROI is excluded, since it can obscure the interpretation of the correlation analysis. For example, *f_sphere_* and MK are both close to 0 in CSF, although they are not necessarily positively correlated in tissue ROIs, thus including CSF would make the correlation analysis more difficult to quantify and interpret.

### 2.6 Comparison with the atlas

The next analysis focused on comparing the parameters estimated from SANDI and DKI with the Allen Brain mouse atlas [68]. Specifically, we investigated how well different dMRI parameters correlate with the image intensity of the atlas generated based on serial two-photon tomography images, in which “dark” and “light” areas often reflect cell density[68].

#### 2.6.1 Atlas information

For this analysis we employed the P56 Mouse Brain atlas which was built at a resolution of 10 μm isotropic using serial two-photon tomography (STPT) images from 1675 young adult mice [68]. In the original STPT images with an in-plane resolution of 0.35μm, the cellular nucleus appears dark and the cytoplasm bright. When interpolating the atlas at 10 μm, the areas with relatively densely packed cells appear bight, for instance layer 4 of the cortex, while areas with sparser cell body distribution appear darker, such as in white matter fibre tracts. Nevertheless, there are some well-known exceptions: for instance the densely packed pyramidal layer of the hippocampus where the large number of nuclei which appear dark in the original STPT images also result in a dark layer in the interpolated image. Thus, to avoid any issues with misregistration when the same gray level intensity reflects very different underlying cell densities, we decided to exclude hippocampal regions from analyses.

#### 2.6.2 Registration

To create the template, we downloaded the P56 Mouse Brain Atlas in NIFTY format with a resolution of 20 μm isotropic (https://scalablebrainatlas.incf.org/mouse/ABA_v3). Then, the atlas was further downsampled to match the resolution of the dMRI data, namely 0.12×0.12×0.4 mm.

To compare the atlas intensity with MRI derived parameters, we performed a 2D, slice to slice registration, to avoid interpolation between MRI slices during the registration process. First, we chose a slice of interest in the atlas, then we picked the corresponding slices from the dMRI dataset based on anatomical landmarks for each animal, and next we performed a 2D non-rigid registration using the *Registration* function from the ANTs package [69] for Python 3 (Python Software Foundation). We tested 2D registrations both using b = 0 ms/μm^2^ images as well as the *f_soma_* maps, and the latter provided a better registration outcome upon inspection, likely because low signal intensity values in the atlas corresponded to some extent to low *f_soma_* values, both in tissue as well as in the ventricles. Thus, the 2D slice registration was performed based on the *f_soma_* maps for all animals and then the transformations were applied to all other parameter maps using a linear interpolation. The registered parameter maps were then averaged over the 6 animals.

#### 2.6.3 Correlation analysis

To study the link between the MRI derived parameters and the image intensity of the atlas, we have considered two representative slices, covering the cerebrum and the cerebellum. Then, for each slice we manually drew an ROI which encompasses the tissue areas, while avoiding the ventricles, for two reasons: a) the MRI data showed larger ventricles compared to the atlas, thus this area was more prone to misregistration b) including ventricles would bias the correlation analysis for parameters such as MD, FA and *f_neurite_* where we would no longer expect a monotonical relationship with atlas intensity. The delineated ROIs are shown in Figure 8. Then, for all voxels in the ROI we computed the Spearman rank correlation coefficient between the dMRI parameters and the atlas intensity, a metric that reflects a monotonic relationship between variables, rather than a linear relationship.

### 2.7 Acceleration

The last analysis of this study investigated the effect of reducing SANDI’s number of shells and maximum b-value, aiming at potentially enabling the acceleration of its acquisition. This was performed by progressively excluding the last shells (higher b-values) from the full protocol employed in this study, which consists of 8 shells with a maximum b-value of 12.5 ms/ μm^2^. Thus, we have analysed data for 7 shells with b = {1, 2.5, 4, 5.5, 7, 8.5, 10} ms/μm^2^, 6 shells with b = {1, 2.5, 4, 5.5, 7, 8.5} ms/μm^2^, and 5 shells with b = {1, 2.5, 4, 5.5, 7} ms/μm^2^. We have not included protocols with less than 5 shells, as this is the number of free parameters estimated by the SANDI model in the current implementation. To perform the model fitting, for each sub-protocol we have retrained the RF regression algorithm on simulated data with the respective number of shells, and the same SNR as detailed in section 2.2.3. Then, real data was employed to estimate the SANDI parameters, which were evaluated voxelwise in the ROIs defined in section 2.5.

## 3. Results

### 3.1 Data quality assessment

To explore the quality of the acquired data, Figure 2 depicts, for a representative mouse, the normalized and directionally averaged raw data, i.e. prior to the application of the denoising, ghost-correction and unringing steps in the pre-processing pipeline. The data is presented for 5 b-values from 0 to 10 ms/μm^2^, both for real and magnitude data, in two slices from a representative animal. The maps show a faster signal attenuation in gray matter compared to white matter and a good contrast between WM and GM at very high b-values (b > 7 ms/μm^2^). Moreover, at these b-values when the signal is highly attenuated (^~^ 5% in GM and ^~^ 15% in WM), we observe most differences between real and magnitude data due to the Rician noise floor, with real data showing lower signal intensity especially in GM and CSF. Similar data quality is observed throughout the brain, as illustrated in Figure S1 in Supplementary Information, which plots the directionally averaged real data for b = 1 and 12.5 ms/μm^2^ and 32 slices in one representative animal. The observed superior-inferior image gradient is due to the sensitivity profile of the receiver cryoprobe, which is a surface coil. This pattern can also be observed in the SNR maps which have lower values in the inferior part. Representative SNR maps are illustrated in Figure S2 in Supplementary Information. The average SNR of the raw data across the entire brain was 37.9 ± 8.2, with higher values in gray matter, for instance SNR = 50.1 ± 12.7 in the cortex, and lower in white matter, for instance SNR = 31.1 ± 5.2 in the internal capsule. After denoising the SNR increased by a factor of ^~^1.3.

### 3.2 SANDI parameter for magnitude and real data

Figure 3 presents the parametric maps derived from SANDI, for real and magnitude data following the pre-processing pipeline described in Figure 1, in one representative animal.

**Figure 3.**
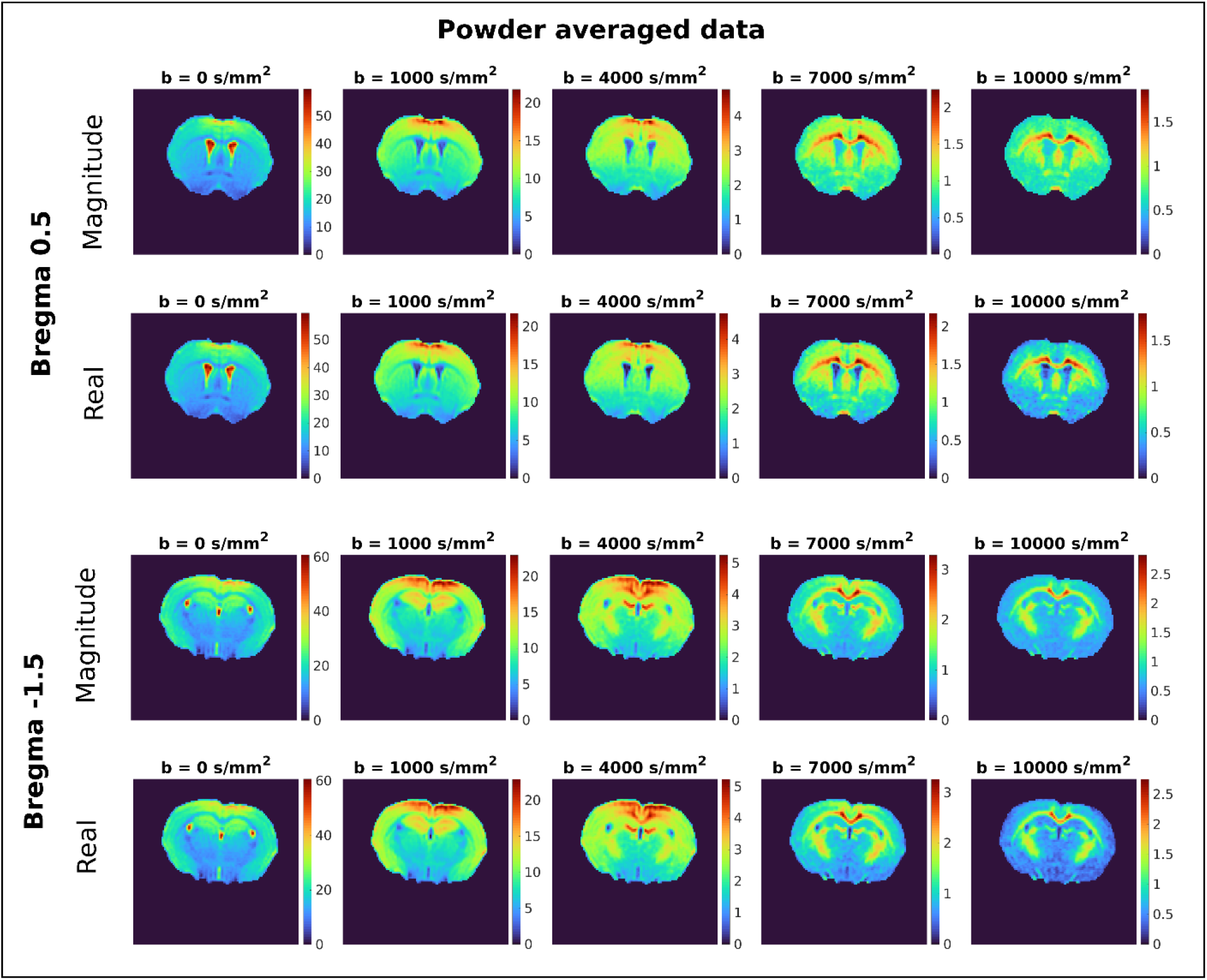
Powder-averaged diffusion weighted images using either magnitude or real data, from one representative animal. The data are presented for two slice positions, and 5 different b values increasing left to right. Clearly, the signal to noise of the powder averaged data is high even at the highest b-value used in this study (b = 10 ms/μm^2^), enabling the downstream processing of the SANDI model.

Qualitatively, a good correspondence between the parametric maps derived from real and magnitude data is observed. Nevertheless, the maps derived from real data exhibit better superior-inferior homogeneity, which can be clearly observed especially in the slice position at Bregma 0.5, where an increase in *f_stick_* and *R_sphere_* and a decrease in *f_sphere_* and *D_stick_* in the inferior part of the brain was observed when parameters were estimated based on magnitude data compared to when they were estimated using real data. Importantly, the coil signal profiles have a decreasing gradient along the superior-inferior axis, as illustrated in Figure 3, with higher signal closer to the surface of the coil, suggesting that Rician bias will be higher in the lower S/N areas in the inferior part of the brain.

We then investigated the more quantitative relationships between real and magnitude signals. The correlation between parameters derived from magnitude and real data in different WM and GM ROIs is shown in Figure 4. Overall, a good correlation between data types was noted, especially for *f_stick_* and *D_ball_* with correlation coefficients r > 0.8. For *f_sphere_* we also see high correlations in WM (r > 0.85), and slightly lower in GM (0.5 < r < 0.8), with lower values in areas where the parameters are more homogenous and have a lower dynamic range, such as striatum (r ^~^ 0.5). For *R_sphere_* the correlation strength is lower, especially in GM (r ^~^ 0.5-0.6). All correlations are significant with p << 0.01. In terms of bias, which is reflected in Figure 5 as a departure from the identity line, *f_stick_* shows the largest differences, with higher values when estimated from magnitude data. *R_sphere_* also shows slightly larger values when estimated from magnitude data, while the other parameters show little bias, as also illustrated in the Bland-Altman diagrams in Supplementary material.

**Figure 4.**
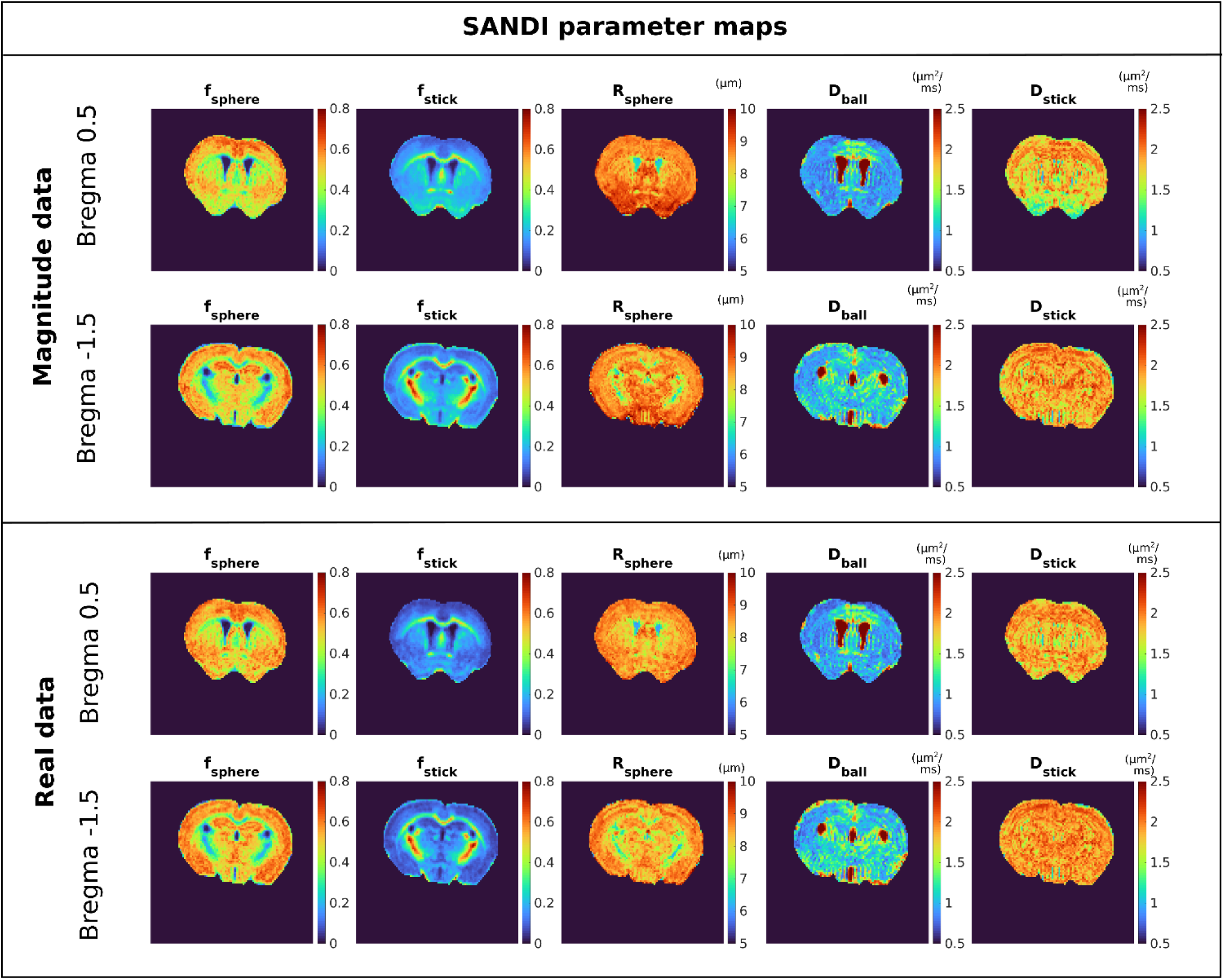
SANDI parameter maps derived from magnitude (top) and real (bottom) data in one representative animal. Note the lower, but non-zero sphere (tentatively assumed to represent cell soma) fraction in white matter, and higher sphere fractions in gray matter. The opposite is observed for the stick fraction (tentatively assumed to represent neurites). In the ventricles, especially in the real data, neither sphere nor stick signals are detected. Sphere radii are quite uniform across the gray matter, and are somewhat lower in white matter regions, while the diffusivity in sticks is between free diffusion (e.g. ventricles in D_ball_) and the other Gaussian diffusion processes in the brain (D_ball_ in the brain parenhcyma), tentatively associated with extracellular diffusion.

**Figure 5.**
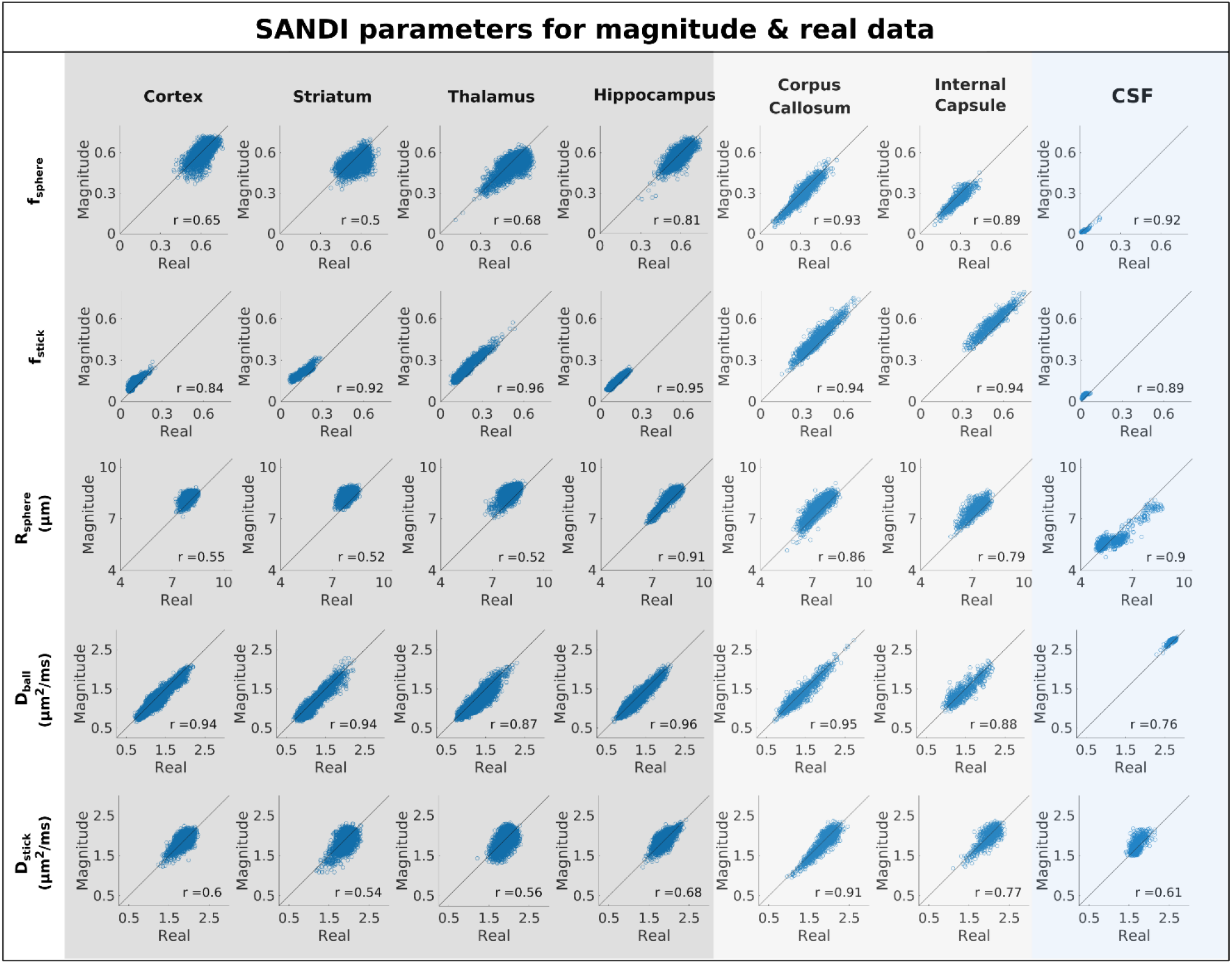
Scatter plots of SANDI parameters estimated based on real and magnitude data for the ROIs defined in Figure 2. The data is shown for GM ROIs (cortex, striatum, thalamus and hippocampus) depicted with dark grey, WM ROIs (corpus callosum and internal capsule) depicted with light grey, and CSF depicted with light blue. While correlations are generally good, some bias is observed in the magnitude data.

For all further analyses, SANDI parameters are estimated from real data, in order to decrease the bias due to Rician noise floor in the powder averaged data [50, 51]. Maps of SANDI parameters for all slices are shown in one representative animal in Figure S1 in Supplementary Information.

### 3.3 SANDI parameter distributions within the ROIs

Next, we analyse the estimated metrics within well-defined ROIs representing similar GM, WM and CSF structures across the animals. Figure 6a shows the decay of the normalized, powder averaged signal in each ROIs as a function of b-value, plotting the data points, the signal prediction from the estimated SANDI parameters, as well as their difference, for one representative animal. Figure 6b presents boxplots of the estimated SANDI parameters from the ROI averaged signals, showing consistent values of the estimated metrics across animals. The results also attest that the model provides a good fit to the data, both when fitted to the ROI averaged signal, as illustrated in Figure 6a, as well as when fitted to individual voxels selected from each ROI, as shown in Figure 6c.

**Figure 6.**
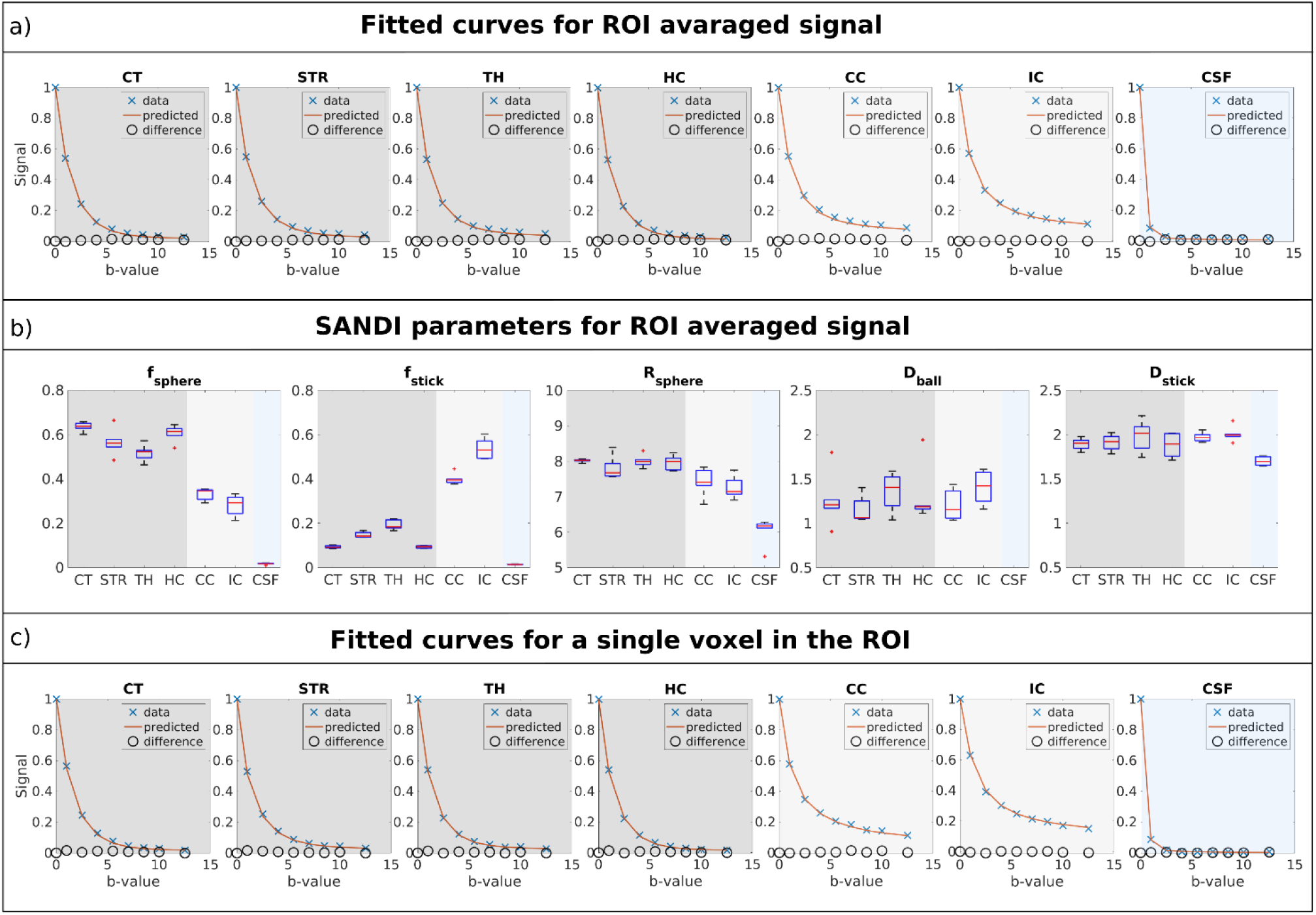
a) Decay curve as a function of b-value of the powder averaged mean ROI data, predicted signal from the estimated SANDI model parameters, as well as their difference in GM, WM and CSF ROIs: CT – cortex, STR – striatum, TH – thalamus, HP – hippocampus, CC – Corpus Callosum, IC – internal capsule, CSF – cerebro-spinal fluid. The curves are presented for one representative animal and show a good fit of the SANDI model to the data. b) Boxplots of estimated SANDI parameters across the six animals in the different ROIs. c) Same as a), but for data from a single voxel selected from each ROIs. Both ROI-based and single-voxel data are deemed to have sufficient signal to noise to be fitted to the SANDI model.

We then turn to investigate the distribution of the metrics within the ROIs. Figure 7 presents histograms of SANDI and DKI parameters in GM, WM and CSF ROIs across the six mice and Table S2 in supplementary material presents the average and standard deviation across animals of the mean parameter values in each ROI. The results reveal several interesting findings. First, higher soma signal fractions were observed in GM than in WM, while the contrast for the neurite signal fraction is reversed with higher values in WM and lower in GM, consistent with previously published results[39–41]. Second, both *f_sphere_* and *f_stick_* are almost 0 in CSF. The other parameters vary less across tissue ROIs, as also illustrated by the mean values in Table S2. In CSF, *D_ball_* approaches the diffusivity of free water. For DKI parameters, MD shows the least contrast between GM and WM, while both FA and MK values are higher in WM compared to GM ROIs.

**Figure 7.**
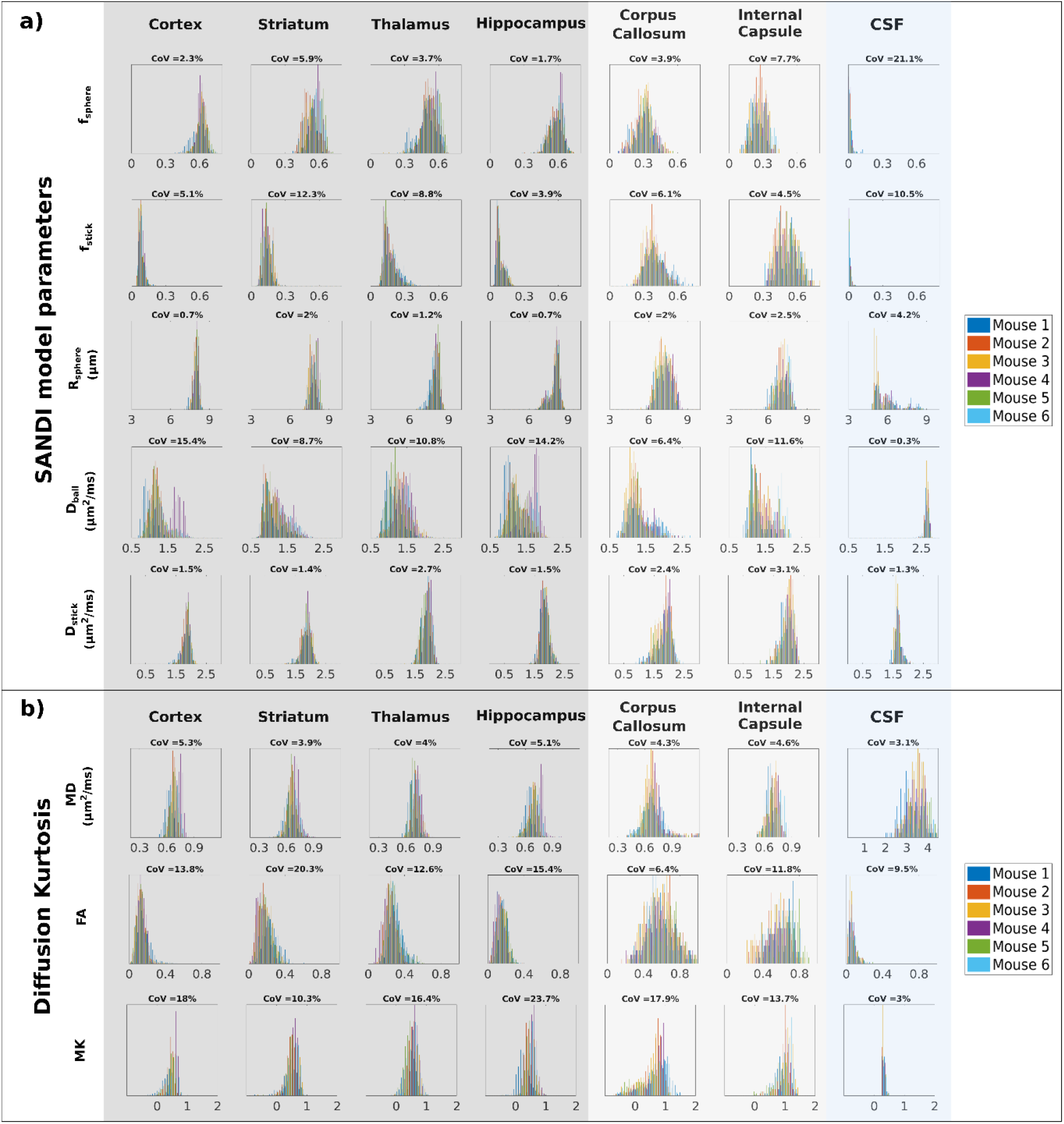
Histograms of a) SANDI and b) DKI parameters for the 7 ROIs depicted in Figure 2. The six animals are presented by different colours. The CoV across animals is calculated for the mean parameter values in each ROI. Excellent stability of the SANDI parameters is observed between the animals.

In terms of variability across animals, we found that *f_sphere_* and *f_stick_* have a coefficient of variation (CoV) up to ^~^12% (range: 1.7% - 12.3%) in tissue ROIs and higher CoVs in CSF due to the very low *f_sphere_* and *f_stick_* mean values. *R_sphere_* and *D_stick_* have a much lower CoV across animals, up to 3.1% in tissue ROIs. *D_ball_* has a higher CoV with values between 6.4% and 15.4%. The variability of *f_sphere_* and *f_stick_* is similar to the variability shown by MD, while FA and MK parameters estimated from the DKI analysis show higher variability across animals (range: 6.4% to 24%).

### 3.4 Do SANDI and DKI parameters correlate well?

The next analysis focused on attempting to establish links between DKI and SANDI parameters. Figure 8 shows scatterplots between DKI and SANDI parameters when the data from all tissue ROIs (i.e. excluding CSF, c.f. Methods) was pooled together.

**Figure 8.**
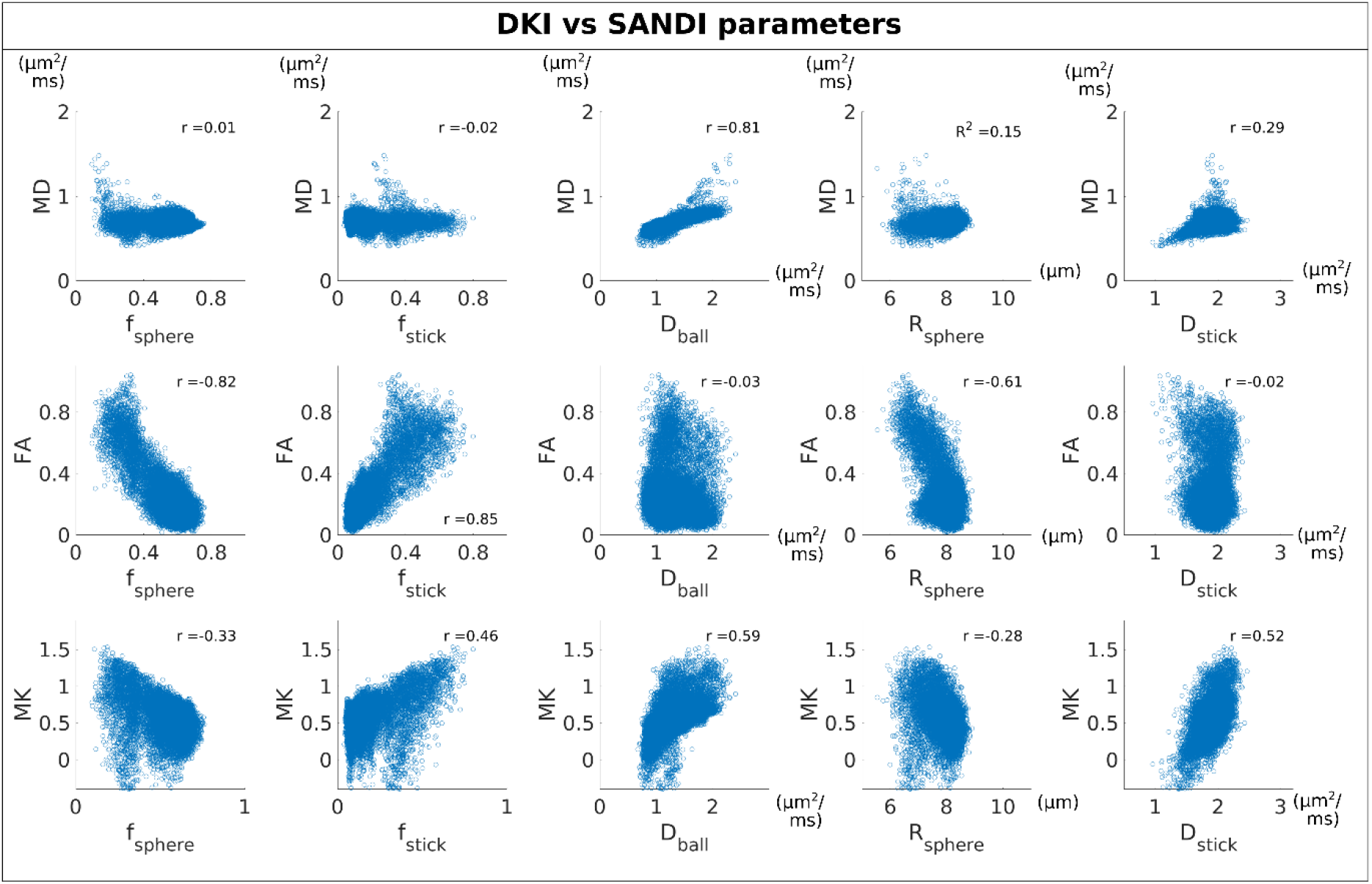
Scatterplots of DKI parameters vs SANDI parameters with datapoints pooled together for all tissue ROIs (excluding CSF). The correlation coefficient is also provided for each plot. MD shows a very strong positive correlation with D_ball_, FA shows a very strong positive correlation with f_sphere_ and negative correlation with f_stick_ and MK shows only moderate and weak correlations with any of the SANDI parameters.

The results show that mean diffusivity has a strong positive corelation with SANDI-derived *D_ball_* (r = 0.8), and a weak positive correlation with *D_stick_* (r = 0.29). The MD did not correlate with the other SANDI parameters (r < 0.15). On the other hand, FA has a very strong positive correlation with *f_stick_* (r = 0.85), a very strong negative correlation with *f_sphere_* (r = −0.82) and a moderate negative correlation with *R_sphere_* (r = −0.61).

MK better correlated with several SANDI parameters, but nevertheless the correlations observed were only moderate. Specifically, we found a positive correlation of MK with *f_stick_, D_ball_* and *D_stick_*(r = 0.46, r = 59, r = 0.52, respectively) and a weak negative correlation with *f_sphere_* and *R_sphere_* (r = −0.33, r = −0.28, respectively).

### 3.5 Comparison of SANDI and DKI parameters with the Allen brain atlas

This analysis is focused on the comparison dMRI parameters derived from SANDI (*f_sphere_* and *f_stick_*) and DKI (MD, FA, MK) with the image intensity of the P56 Allen brain atlas, which reflects cell density (among other things) [68]. Figure 9 show the results for two representative slices covering both the cerebrum (a) and the cerebellum (b).

**Figure 9.**
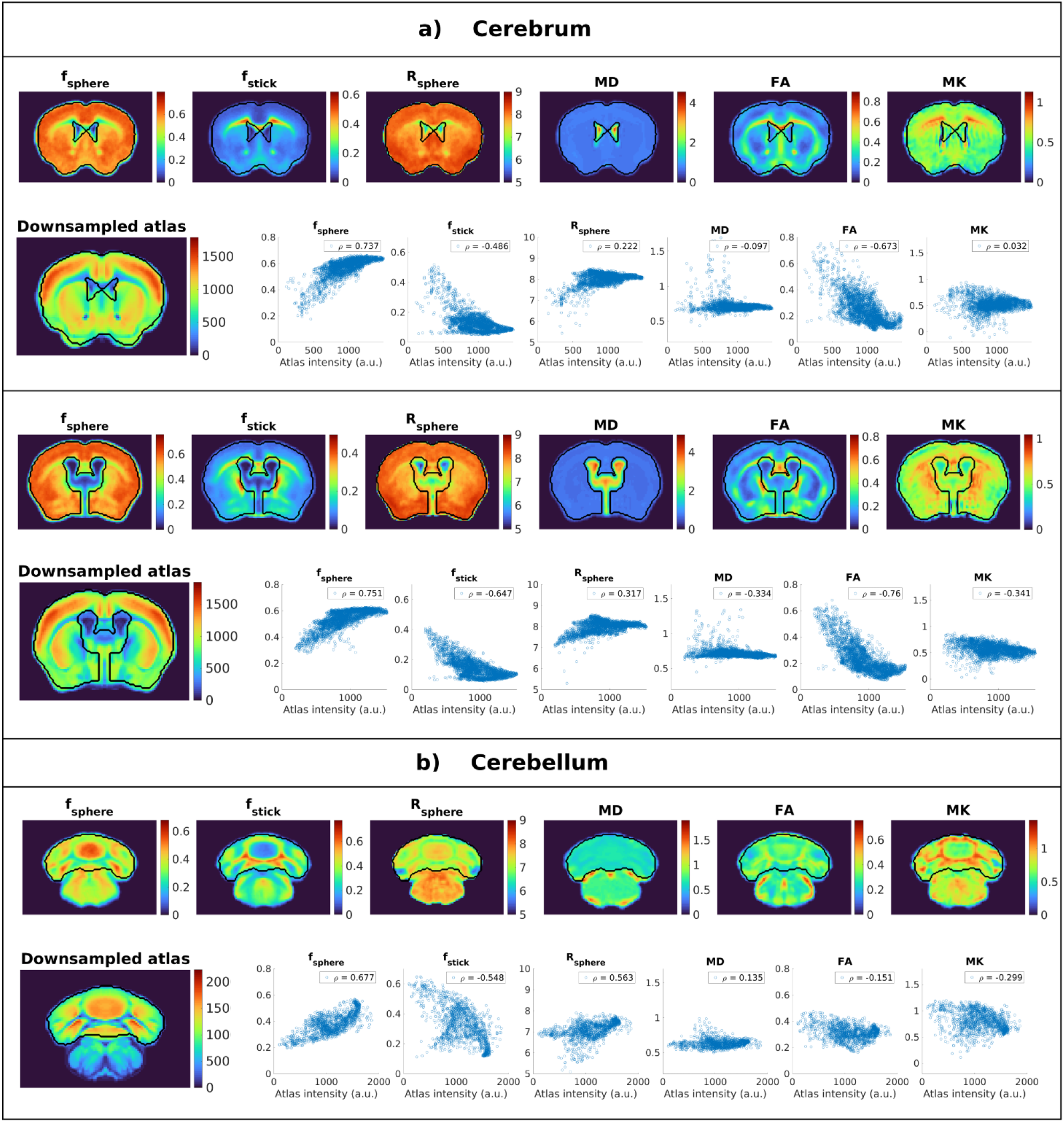
Comparison between dMRI parameters and the downsampled Allen mouse brain atlas for three slices in the a) cerebrum and b) cerebellum. For each region, the top row shows the parameter maps derived from SANDI (f_sphere_, f_stick_ and R_sphere_) and DKI (MD, FA and MK), averaged over the 6 animals, after registration to the template as described in section 2.6.2. The bottom row shows voxelwise scatter plots of the same dMRI parameters versus the image intensity of the atlas, and the legend presents the Spearman correlation coefficient. Strikingly, f_sphere_ exhibits a very good rank correlation with the Allen brain contrast in all areas.

Both in the cerebrum and cerebellum, the maps of the group *f_sphere_* qualitatively follow the intensity patterns observed in the downsampled Allen atlas, with higher values in the gray matter and lower in the white matter. Although group *f_sphere_* does not exhibit the same dynamic range as the atlas image intensity in gray matter, it does provide contrast between different regions, for example it has lower values in striatum compared to the cortex. The scatterplots also show there is a strong positive rank correlation between group *f_sphere_* and the atlas intensity with Spearman correlation coefficients of ρ = 0.74, ρ = 0.75 and ρ = 0.68, respectively.

In the cerebrum, a strong negative correlation is also observed between the atlas intensity and FA (ρ = −0.67 and ρ = −0.73), and a moderate negative correlation with *f_stick_* (ρ = −0.49 and ρ = - 0.64). For *R_sphere_*, MD and MK, we see only weak and very weak correlations (ρ = 0.22 and ρ = 31 for *R_sphere_*; ρ = −0.1 and ρ = −0.33 for MD; ρ = 0.03 (n.s.) and ρ = −0.34).

In the cerebellum, the only parameter which shows a strong correlation with the atlas intensity is *f_sphere_*, nevertheless, we also see moderate correlation with *f_stick_* and *R_sphere_*, with ρ = −0.54 and ρ = 0.56, respectively. For DKI parameters we see weaker correlations, specifically, ρ = 0.14 for MD, ρ = −0.15 for FA and ρ = −0.3 for MK. All correlation coefficients are significant with p << 0.01 unless otherwise specified.

Although this analysis has been performed on the group averaged metrics, similar patterns were observed in individual mice, as illustrated in Figure S6 from the Supplementary material for the SANDI parameters.

### 3.7 Accelerating SANDI acquisitions

Finally, we investigate the potential for accelerating the acquisitions by removing data. Figure 10 illustrates the SANDI parameters estimated from protocols with decreasing number of shells for the different GM, WM and CSF ROIs considered in this work. The results show that the estimated parameter values are overall stable when decreasing the acquisition protocol to 5 shells, both in terms of median values and interquartile ranges, which reflect variability within the ROIs as well as between animals. In most cases presented in Figure 9, the median values estimated from the 5-shell protocol are within 10% of the values estimated from the full protocol, as shown by the dotted green lines. In other cases, there is a small change in parameter values, for instance, an increase in the estimated *R_sphere_*, which is more pronounced in the WM ROIs, as well as an increase in *f_stick_* in the GM ROIs. For *R_sphere_*, the median values are still within 10% difference, while for *f_stick_* the absolute differences are small, i.e. median value differences < 0.03, although in GM this change is larger than 10% due to the overall small values of the parameters.

**Figure 10.**
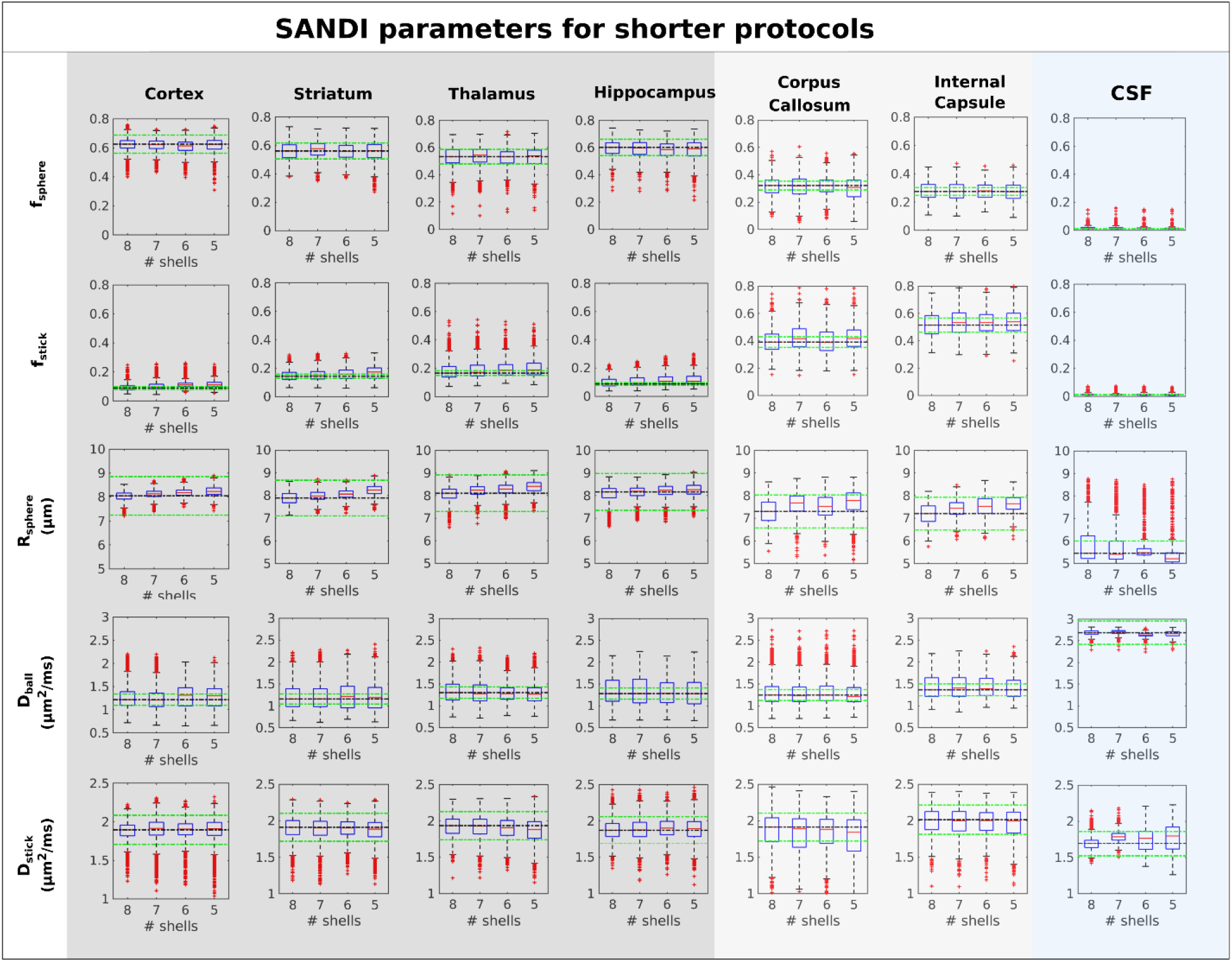
Boxplots of estimated SANDI parameters for protocols with varying number of shells in GM, WM, and CSF ROIs. The parameters are estimated voxelwise and aggregated over different animals for the voxels in each ROI. The rows present different model parameters and the columns different ROIs. The dotted black lines show the values estimated from the full, 8-shell, protocol, while the green lines show a variation of ±10 % from this value. The b-values employed in the different protocols are the following: full protocol (8 shells): b = {1,2.5,4,5.5,7,8.5,10,12.5} ms/μm^2^; 7-shells: b = {1,2.5,4,5.5,7,8.5,10} ms/μm^2^; 6-shells: b = {1,2.5,4,5.5,7,8.5} ms/μm^2^ and 5-shells: b = {1,2.5,4,5.5,7} ms/μm^2^. Overall, all SANDI parameters are stable as the protocol is reduced from 8 shells with maximum b-value of 12.5 ms/μm^2^ to 5 shells with maximum b-value of 7 ms/μm^2^.

## 4. Discussion

Preclinical animal models are heavily utilised to investigate many facets of normal biological process such as development[70], aging[71, plasticity, learning and memory, 72], as well as adverse effects such as neurodegeneration [73], stroke [74] and other brain injuries [75], and cancer [76]. Due to their non-invasive nature and ability to probe microscopic dimensions, diffusion MRI (dMRI) methods have been particularly useful for reporting on progressive changes in tissue microstructure, for instance in the detection of Alzheimer’s disease [77], demyelination [78], etc. However, most dMRI methods focused on WM due to the somewhat more simplistic nature of the structures (at least in terms of tissue modelling). By contrast, the SANDI model was designed to capture features that can be more relevant in gray matter, thereby providing an exciting opportunity to enhance microstructural inference via dMRI measurements.

Here, we implemented SANDI on a 9.4T pre-clinical scanner and characterized, for the first time, the mouse brain microstructure, in-vivo, and attempted to correlate the SANDI findings with the Allen mouse brain atlas to provide a first “group-level” validation of the technique. Our work also suggests important magnitude versus real data effects and optimisations of the processing pipeline for in vivo imaging. Finally, we examined the relationship between SANDI and more standard DKI parameters as well as the possibility of reducing the acquisition protocol. All these as well as limitations and potential future implications, are discussed below.

### SANDI parameters across regions and animals

Overall, we find that in the in-vivo mouse brain, SANDI acquisitions produce parameters trends that are in line with previous studies performed in vivo in humans [30, 39–41] (and one case of ex vivo mouse brain [39]), with higher *f_sphere_* values in GM than in WM, while the trends for *f_stick_* are reversed. In CSF, both metrics are close to zero, providing an important internal reference that strong biases are not present where very few cells are expected. This can be important for investigating diseases where cell density decreases, for instance due to neuronal loss in Alzheimer’s [79] or Parkinson’s [80] disease. More specifically in GM ROIs, we saw a narrow distribution of *f_sphere_* values with a mean of 0.59±0.02 in the cortex, while in the deeper GM regions and hippocampus, the distributions were wider, reflecting the more heterogenous sub-structures within these areas [81]. Although previous ex-vivo SANDI data which was acquired with much higher resolution (0.05 × 0.05 × 0.25 mm) showed clear differences between cortical layers [30], here such differences are not obvious, likely due to the lower resolution, both in plane as well as slice thickness, namely 0.12 × 0.12 × 0.4 mm in our study.

In WM, as expected, SANDI produces lower estimates of *f_sphere_* values (^~^0.3) and the distributions are wider than in GM, also due to larger partial volume effects considering the (still very thin) slice thickness of 0.4 mm employed in this *in vivo* acquisition. Importantly, many models simply ignore spherical objects (assumingly, cell bodies) in white matter modelling of dMRI. Indeed, histological findings suggest that cell body density (not only neurons, but also astrocytes, microglia, etc) in the white matter is certainly not negligible [30, 82]. Our results also show that tissue components which are characterized by slow isotropic diffusion contribute to the signal with a fraction ^~^0.3 in WM, consistent with previous results [39, 83], and should not be ignored. Overall, the coefficients of variation of the mean *f_sphere_* values across animals were small, < 8% both in GM and WM, showing a good reproducibility of the metric.

As some of its predecessors[22–24, 31, 84], SANDI also provides parametric estimates for the fraction of neurites, which are modelled as sticks, *f_stick_*. Using SANDI, the *f_stick_* metric showed lower values in GM, between 0.1 in cortex and 0.2 in thalamus, and higher values in WM (0.39±0.03 in CC and 0.52±0.02 in IC). The histograms were also narrower in cortex and wider in WM, due to partial volume effects, and in deep GM, especially in the thalamus where there are more myelinated fibres compared to the other GM ROIs considered here. The coefficient of variation for *f_stick_* is also relatively small < 12%. The patterns and values of *f_stick_* measured in this study are in line with previous works estimating the signal fraction of sticks, both for techniques which use directional data [22–24, 84], as well as powder averaged data [39, 40] [30, 41]. In CSF, both *f_sphere_* and *f_stick_* are close to zero with a narrow distribution of values, although the CV is higher due to the division by the low mean values.

*R_sphere_* values also show contrast between brain regions, with lower values in WM than in GM. Overall, the estimated *R_sphere_* values are between 6 and 9 μm, consistent with values derived from histology, when considering that the apparent effective radii are actually tail-weighted values of the underlying distribution of sizes [85]. Higher *R_sphere_* values can also be observed in regions known to have larger cells, such as the granular and pyramidal layer of the hippocampus or the piriform cortex[81], as illustrated in Figure S7 in Supplementary Information. The choice of sequence parameters, especially the gradient duration and diffusion time, can also play a role on the sensitivity to different cellular sizes and might influence the estimated *R_sphere_* values.

The other SANDI parameters, namely *D_ball_* and *D_stick_* are also reproducible across animals, although the coefficients of variation for *D_ball_* are higher than for *D_stick_* with values of 6-16% and <4%, respectively. The larger CoV of *D_ball_* is driven mostly by the values from one animal which are higher, and also visible as a separate peak on the MD histograms. Overall, these parameters show the least contrast across tissue types, and they seem to capture the effect of imaging artifacts, for instance Gibbs ringing due to Partial Fourier, as can be seen in Figure 4, where some ringing around ventricles can be seen in the maps of *D_stick_* and *D_ball_*, but not for the other parameters.

### Comparison of real and magnitude data

Our findings suggest that real data is preferable for more accurate SANDI estimation, at least in cases where signal to noise is similar or poorer than in this study. Comparing the parameters estimated from magnitude and real data, the Rician noise floor clearly biases SANDI-derived parameters, mostly *f_stick_*, which has significantly higher values when magnitude data is considered. Still at these levels of signal to noise, there is a strong positive correlation between the two cases, especially in ROIs with a larger dynamic range of parameter values where r > 0.9. For data with lower SNR, for instance on clinical scanners, better averaging approaches can also be employed to reduce the Rician noise bias compared to taking the mean over directions [86]. Other parameters show less bias, although we observe a small positive bias for *R_sphere_* and a small negative bias for *f_sphere_*, which is consistent with a previous simulation study which shows that in the presence of Rician noise, *R_sphere_* tends to be overestimated, especially for lower values of *f_sphere_*, such as in WM ROIs[40].

### Comparison between DKI and SANDI parameters

We sought to compare DKI and SANDI parameters to establish whether perhaps performing DKI can be a good proxy for SANDI. The results revealed that MD is strongly correlated to SANDI’s *D_ball_*, FA is strongly correlated to SANDI’s *f_sphere_* and *f_stick_* and MK is only moderately correlated to any of the SANDI parameters. Although FA strongly correlates with SANDI’s critical *f_sphere_* parameter for the ROIs considered here, it does not correlate as well with the Allen brain atlas, especially in the cerebellum (r = −0.16), suggesting SANDI cannot be merely replaced by DKI.

Another point is the variation of parameters due to (imaging) artifacts. While in SANDI, the local variation in image intensity, for instance due to Gibbs ringing, is mostly reflected by the *D_ball_* and *D_stick_* parameters, as illustrated in Figure 4, for DKI such intensity variations are reflected especially in the kurtosis maps, also giving rise (as expected) to the negative kurtosis values[87] seen in Figure 7 and 8. Such effects are present especially in the corpus callosum as it is located next to the ventricle, an area of large image contrast thus prone to Gibbs ringing artifacts, especially that data was acquired with Partial Fourier, which were partially, but not entirely removed during pre-processing. These effects on MK are consistent with previous literature, for instance the maps presented in [78, 88], where there are obvious drops in MK values due to Gibbs ringing.

Moreover, the DKI analysis included a subset of the measurements with only two b-shells of {1, 2.5} ms/μm^2^ which are in the range of diffusion weightings appropriate for DKI [65], resulting in more variability of the maps compared to the SANDI parameters which were derived from 4x more measurements. Although, DKI requires less b-values and a faster acquisition, it appears that the high b-values used in the SANDI approach do provide additional contrast, as discussed below.

### Comparison to the Atlas

The results from this study show a strong positive rank correlation between SANDI-driven *f_sphere_* and the Allen brain atlas image intensity, which reflects to some extent cell density [68]. *f_sphere_* is the only parameter which has a strong correlation both in the cerebrum and cerebellum, while other parameters such as FA and *f_stick_* show a strong / medium correlation only in the cerebrum, but not the cerebellum. These results suggest that there is information in the metrics derived high diffusion weighted data, which cannot be captured by DTI / DKI parameters. Another possible explanation is the effect of the slice thickness in the dMRI data. While in the cerebrum, there is a smoother variation in the tissue with changes in the slice direction, the cerebellum is smaller, and the tissue architecture changes more over the 0.4 mm slice thickness. This especially affects metrics such as FA which will have lower values where there are larger intra-voxel variations in fibre direction. Other possible issues are related to the non-rigid registration, as quite large deformations are necessary to match the EPI based dMRI images to the atlas due to susceptibility artifacts.

Moreover, the registration was performed based on *f_sphere_* maps, which visually provided a better match between the atlas and the dMRI data compared to using b = 0 ms/μm^2^ images, either directly registered to the atlas, or registered first to anatomical undistorted images and then to the atlas. Although this choice might favour the correlation between *f_sphere_* and the atlas intensity, very similar patterns to the average map can be observed in individual maps before registration, as shown in Figure S3 in Supplementary material for all 6 mice, visually attesting that areas of high image intensity in the atlas correspond to those with high *f_sphere_* values.

Although we see a strong rank correlation between *f_sphere_* and the Allen brain atlas, the atlas shows a wider dynamic range of intensity values compared to *f_sphere_*, especially in the cortex. This is likely related to the different contrast mechanisms of the two imaging modalities. In the two-photon tomography employed for the Allen Brain Atlas, the image intensity is related to the amount of cytoplasm, while *f_sphere_* reflects the amount of dMRI signal coming from water molecules restricted in spherical environments. While both metrics are considered to reflect to some extent cell density their contrast is inherently different, so we would not necessarily expect a perfect correlation. One example is the hippocampus, where the pyramidal layer has highly packed cells, as illustrated by stains such as Nissl or DAPI. In the Allen brain atlas, the pyramidal layer appears dark, due to the large and densely packed cell nuclei, nevertheless, *f_sphere_* values are high in this region.

### Acceleration of SANDI

The last analysis of this study investigated the possibility to shorten the SANDI acquisition time as well as reduce the maximum b-value, which would facilitate lower echo times, better SNR, as well as acceleration that favours clinical application on standard systems with limited gradient strengths. In the past SANDI was performed with data from high performance clinical scanners, as well as standard hardware by combining linear and spherical tensor encoding. Our results show that metrics derived from standard PGSE acquisitions, which are widely available on clinical and pre-clinical systems, are comparable with previous values from the literature and are overall stable in both WM and GM ROIs, when the protocol is reduced from 8 shells with a maximum b-values of 12.5 ms/m^2^ to 5 shells with a b-value of 7000 s/mm^2^, which is very encouraging for future applications.

### Limitations

As any study, ours also has some limitations, both intrinsic to the SANDI model, as well as related to the experiments.

The SANDI model assumes non-exchanging compartments, where cell bodies are represented as spheres with apparent radius *R_sphere_*, neurites (axons and dendrites) are represented as sticks, and the extracellular space is assumed to be a well-mixed environment characterized by Gaussian diffusion. This is a simplified view of the tissue and does not account for its full complexity, for instance variations in cell sizes and shapes, finite axon and dendrite sizes, the presence of intracellular organelles, membrane permeability and exchange across different compartments, etc. In terms of diffusion modelling, within the GPD approximation, the kurtosis of each compartment is assumed negligible. As recent studies employing double diffusion encoding have shown non-zero intra-compartment kurtosis in brain tissue both using clinical and pre-clinical data [89–91], these assumptions might also have an impact on the estimated parameter values, and their interpretation. Another modelling assumption is the lack of exchange between compartments. According to some studies, the intra-cellular residence time of water is on the order of 500 ms [92], which would not play an important role at the diffusion time considered here (20 ms), although other recent studies show that water exchange across compartments can be faster in GM and can influence the dMRI signal at this diffusion time [93]. PGSE data acquired at a single diffusion time, as done in this study, is not appropriate to characterize the effects of intra-compartment kurtosis or exchange, which might produce some biases in the estimated parameters, that can be the focus of future work.

In terms of Atlas comparison, the SANDI metrics and the Atlas image intensity, which are used as proxies for cell density in the brain, are based intrinsically on different contrast mechanisms and assumptions, thus we do not expect to see a perfect correlation. Moreover, the dMRI metrics are compared to an average template, rather than data specific for each animal, and this might incur larger deformations than when comparing to slices from the same brain. Moreover, the template is based on young adult mice that are 8 weeks old, while the animals in our study are significantly older (34 ± 5 weeks), thus the effect of age on brain (micro)structure is not considered. An example of this effect is the area around the ventricles, which is prone to more severe misregistration, as illustrated in Figure 8, since the animals in our cohort have larger ventricles compared to the template. Thus, to avoid bias due to misregistration, these areas were not included in the correlation analysis. Future studies should engage in active histology of the imaged animals and find perhaps better proxies for cell body density than the intensity of the two-photon tomography used in the Allen brain atlas.

In terms of shortening the acquisition protocol, this study has focused on subsets of the current dataset by decreasing the maximum b-value, which, as discussed, has several advantages in practice. Nevertheless, a lower number of shells with other b-value combinations, or even optimising the b-values, the diffusion time and/or the number of directions per shell is likely to further improve the performance of shorter protocol, which is also a possible direction for future research. Moreover, in this study we employed standard PGSE measurements (i.e. linear encoding) with a range of b-values up to very high diffusion weighting (12,5 ms/μm^2^), as described in the original SANDI methodology. Using more advanced diffusion sequences and combining linear encoding with other shapes of the b-tensor, such as planar or spherical encoding, might benefit the estimation of SANDI parameters, although the results of the two different approaches have not been directly compared. Nevertheless, the benefit of using standard PGSE sequences is their wide availability on clinical and pre-clinical systems.

## 5. Conclusions

We characterised mouse brain microstructure using SANDI in vivo and showed that the fraction of spheres – tentatively associated with cell bodies – correlate well with the Allen brain atlas contrast. SANDI shows excellent reproducibility across animals and high parametric stability, and the acquisition protocol can be further shortened to 5 shells without significantly biasing the parameter estimates, which suggests potential clinical translation. These findings bode well for further validation of SANDI vis-à-vis its underlying biological underpinnings and pave the way for future characterisations of gray matter in animal models of disease, plasticity, and development.

## Acknowledgements

This study was funded by the European Research Council (ERC) under the European Union’s Horizon 2020 research and innovation programme (Starting Grant, agreement No. 679058). The authors acknowledge the vivarium of the Champalimaud Centre for the Unknown, a facility of CONGENTO financed by Lisboa Regional Operational Programme (Lisboa 2020), project LISBOA01-0145-FEDER-022170. AI and RC are supported by ”la Caixa” Foundation (ID 100010434) and from the European Union’s Horizon 2020 research and innovation programme under the Marie Skłodowska-Curie grant agreement No 847648, fellowship code CF/BQ/PI20/11760029. JC is supported by ERC (Starting Grant, agreement No. 679058) and by European Union’s Horizon 2020 research and innovation programme under the Marie Sklodowska-Curie grant agreement No 101032056. MP is supported by UKRI Future Leaders Fellowship grant no. MR/T020296/1.

## References

1. Alexander, D.C., et al., Imaging brain microstructure with diffusion MRI: practicality and applications.NMR Biomed, 2017. 32(4): p. e3841.

2. Novikov, D.S., et al., Quantifying brain microstructure with diffusion MRI: Theory and parameter estimation. NMR Biomed, 2019. 32(4): p. e3998.

3. Cassey, B.J., et al., Imaging the developing brain: what have we learned about cognitive development? TRENDS in Cognitive Sciences, 2005. 9(3).

4. Zatorre, R.J., R. Douglas Fields, and H. Johansen-Berg, Plasticity in gray and white: neuroimaging changes in brain structure during learning. Nature Neuroscience, 2012. 15: p. 528–536.

5. Uylings, H.B. and J.M. de Brabander, Neuronal changes in normal human aging and Alzheimer’s disease. Brain Cogn, 2002. 49(3): p. 268–76.

6. Wegiel, J., et al., The neuropathology of autism: defects of neurogenesis and neuronal migration, and dysplastic changes. Acta Neuropathol, 2010. 119(6): p. 755–70.

7. Po, C., et al., White matter reorganization and functional response after focal cerebral ischemia in the rat. PLoS One, 2012. 7(9): p. e45629.

8. Rudrapatna, S.U., et al., Can Diffusion Kurtosis Imaging Improve the Sensitivity and Specificity of Detecting Microstructural Alterations in Brain Tissue Chronically After Experimental Stroke? Comparisons With Diffusion Tensor Imaging and Histology. NeuroImage, 2014. 97: p. 363–73.

9. Shemesh, N., et al., Metabolic properties in stroked rats revealed by relaxation-enhanced magnetic resonance spectroscopy at ultrahigh fields. Nature Communications, 2014. 5: p. 4958.

10. Budde, M.D., et al., Primary Blast Traumatic Brain Injury in the Rat: Relating Diffusion Tensor Imaging and Behavior. Frontiers in Neurology, 2013. 4(154).

11. Reynaud, O., Time-Dependent Diffusion MRI in Cancer: Tissue Modeling and Applications. Frontiers in Physics, 2017. 5(58).

12. Stejskal, E.O. and J.E. Tanner, Spin Diffusion Measurements: Spin Echoes in the Presence of a Time-Dependent Field Gradient Journal of Chemical Physics, 1965. 42: p. 288-+.

13. Novikov, D.S., V.G. Kiselev, and S.N. Jespersen, On modeling. Magn Reson Med, 2018. 79(6): p. 3172–3193.

14. De Luca, A., et al., On the generalizability of diffusion MRI signal representations across acquisition parameters, sequences and tissue types: chronicles of the MEMENTO challenge. bioRxiv, 2021: p. 2021.03.02.433228.

15. Basser, P.J., J. Mattiello, and D. LeBihan, MR diffusion tensor spectroscopy and imaging. Biophysical Journal, 1994. 66(1): p. 259–267.

16. Jensen, J.H., et al., Diffusional kurtosis imaging: the quantification of non-gaussian water diffusion by means of magnetic resonance imaging. Magn Reson Med, 2005. 53: p. 1432–40.

17. Henriques, R.N., S.N. Jespersen, and N. Shemesh, Microscopic anisotropy misestimation in spherical-mean single diffusion encoding MRI. Magn Reson Med, 2019. 81(5): p. 3245–3261.

18. Jelescu, I.O., et al., One diffusion acquisition and different white matter models: how does microstructure change in human early development based on WMTI and NODDI? Neuroimage, 2014. 107: p. 242–56.

19. Jelescu, I.E., et al., Challenges for biophysical modeling of microstructure. Journal of Neuroscience Methods 2020. 108861.

20. Ghosh, A., A. Ianus, and D.C. Alexander, Advanced Diffusion Models, in Quantitative MRI of the Brain, Principles of Physical Measurement, Second edition M. Cercignani, N.G. Dowell, and T.P. S., Editors. 2018.

21. Panagiotaki, E., et al., Compartment models of the diffusion MR signal in brain white matter: a taxonomy and comparison. Neuroimage, 2012. 59(3): p. 2241–54.

22. Fieremans, E., et al., Novel white matter tract integrity metrics sensitive to Alzheimer disease progression. AJNR Am J Neuroradiol, 2013. 34(11): p. 2105–12.

23. Jespersen, S.N., et al., Neurite density from magnetic resonance diffusion measurements at ultrahigh field: comparison with light microscopy and electron microscopy. Neuroimage, 2010. 49(1): p. 205–16.

24. Jespersen, S.N., et al., Modeling Dendrite Density From Magnetic Resonance Diffusion Measurements. NeuroImage, 2007. 34(4): p. 1473–86.

25. Jelescu, I.O., et al., In vivo quantification of demyelination and recovery using compartment-specific diffusion MRI metrics validated by electron microscopy. Neuroimage, 2016. 132: p. 104–14.

26. Panagiotaki, E., et al., Noninvasive quantification of solid tumor microstructure using VERDICT MRI.Cancer Res, 2014. 74: p. 1902–12.

27. Xu, J., et al., Mapping mean axon diameter and axonal volume fraction by MRI using temporal diffusion spectroscopy. Neuroimage, 2014. 103C: p. 10–19.

28. Roberts, T.A., et al., Noninvasive diffusion magnetic resonance imaging of brain tumour cell size for the early detection of therapeutic response. Scientific reports, 2020. 10(1): p. 9223–9223.

29. Kakkar, L.S., et al., Low frequency oscillating gradient spin-echo sequences improve sensitivity to axon diameter: An experimental study in viable nerve tissue. NeuroImage, 2018. 182: p. 314–328.

30. Palombo, M., et al. Histological validation of the brain cell body imaging with diffusion MRI at ultrahigh field. in Proceedings of the ISMRM Annual Meeting. 2019. Montreal, Canada.

31. Behrens, T.E., et al., Probabilistic diffusion tractography with multiple fibre orientations: What can we gain? Neuroimage, 2007. 34(1): p. 144–55.

32. Sotiropoulos, S.N., T.E. Behrens, and S. Jbabdi, Ball and rackets: Inferring fiber fanning from diffusion-weighted MRI. Neuroimage, 2012. 60(2): p. 1412–25.

33. Kaden, E., et al., Multi-compartment microscopic diffusion imaging. Neuroimage, 2016. 139: p. 346–359.

34. Ferizi, U., et al., A ranking of diffusion MRI compartment models with in vivo human brain data.Magn Reson Med, 2014. 72(6): p. 1785–92.

35. Veraart, J., et al., Noninvasive quantification of axon radii using diffusion MRI. eLife, 2020. 9: p. e49855.

36. Olesen, J.L., et al., Beyond the diffusion standard model in fixed rat spinal cord with combined linear and planar encoding. Neuroimage, 2021. 231: p. 117849.

37. Palombo, M., et al. Abundance of cell bodies can explain the stick model’s failure in grey matter at high bvalue. in Joint Annual Meeting ISMRM-ESMRMB. 2018. Paris, France.

38. Stanisz, G.J., et al., An analytical model of restricted diffusion in bovine optic nerve. Magnetic Resonance in Medicine, 1997. 37: p. 103–111.

39. Palombo, M., et al., SANDI: A compartment-based model for non-invasive apparent soma and neurite imaging by diffusion MRI. NeuroImage, 2020. 215: p. 116835.

40. Afzali, M., et al., SPHERIOUSLY? The challenges of estimating sphere radius non-invasively in the human brain from diffusion MRI. Neuroimage, 2021. 237: p. 118183.

41. Gyori, N.G., et al., On the potential for mapping apparent neural soma density via a clinically viable diffusion MRI protocol. NeuroImage, 2021. 239: p. 118303.

42. Palombo, M., et al., Corrigendum to “SANDI: A compartment-based model for non-invasive apparent soma and neurite imaging by diffusion MRI” [Neuroimage 215 (2020), 116835]. Neuroimage, 2021. 226: p. 117612.

43. Ianus, A., et al., Mapping complex cell morphology in the grey matter with double diffusion encoding MRI: a simulation study. arXiv, 2020: p. 2009.11778.

44. Veraart, J., et al., Denoising of diffusion MRI using random matrix theory. Neuroimage, 2016. 142: p. 394–406.

45. Does, M.D., et al., Evaluation of principal component analysis image denoising on multi-exponential MRI relaxometry. Magnetic Resonance in Medicine, 2019. 81(6): p. 3503–3514.

46. Buonocore, M.H. and L. Gao, Ghost artifact reduction for echo planar imaging using image phase correction. Magn Reson Med, 1997. 38(1): p. 89–100.

47. Walsh, D.O., A.F. Gmitro, and M.W. Marcellin, Adaptive reconstruction of phased array MR imagery.Magn Reson Med, 2000. 43(5): p. 682–90.

48. Griswold, M., et al. The Use of an Adaptive Reconstruction for Array Coil Sensitivity Mapping and Intensity Normalization. in Annual Meeting of the ISMRM. 2002.

49. https://github.com/andyschwarzl/gpuNUFFT/blob/master/matlab/demo/utils/adapt_array_2d.m.

50. Fan, Q., et al., Axon diameter index estimation independent of fiber orientation distribution using high-gradient diffusion MRI. NeuroImage, 2020. 222: p. 117197.

51. Eichner, C., et al., Real diffusion-weighted MRI enabling true signal averaging and increased diffusion contrast. NeuroImage, 2015. 122: p. 373–384.

52. Kellner, E., et al., Gibbs-ringing artifact removal based on local subvoxel-shifts. Magnetic Resonance in Medicine, 2016. 76: p. 1574–1581.

53. Motta, A., et al., Dense connectomic reconstruction in layer 4 of the somatosensory cortex. Science, 2019. 366(6469): p. eaay3134.

54. Callaghan, P.T., K.W. Jolley, and J. Lelievre, Diffusion of Water in the Endosperm Tissue of Wheat Grains as Studied by Pulsed Field Gradient Nuclear Magnetic Resonance. Biophys J, 1979. 28(1): p. 133–41.

55. Celebre, G., L. Coppola, and G.A. Ranieri, Water Self-Diffusion in Lyotropic Systems by Simulation of Pulsed Field Gradient-Spin Echo Nuclear-Magnetic-Resonance Experiments. Journal of Chemical Physics, 1992. 97(10): p. 7781–7785.

56. Price, W.S., Pulsed-field gradient nuclear magnetic resonance as a tool for studying translational diffusion. 1. Basic theory. Concepts in Magnetic Resonance, 1997. 9(5): p. 299–336.

57. Kaden, E., F. Kruggel, and D.C. Alexander, Quantitative mapping of the per-axon diffusion coefficients in brain white matter. Magn Reson Med, 2016. 75(4): p. 1752–63.

58. Kroenke, C.D., J.J. Ackerman, and D.A. Yablonskiy, On the nature of the NAA diffusion attenuated MR signal in the central nervous system. Magn Reson Med, 2004. 52(5): p. 1052–9.

59. Jespersen, S.N., et al., Modeling dendrite density from magnetic resonance diffusion measurements.Neuroimage, 2007. 34(4): p. 1473–86.

60. Neuman, C.H., Spin echo of spins diffusing in a bounded medium. J Chem Phys, 1974. 60(11): p. 4508–4511.

61. Balinov, B., et al., The NMR Self-Diffusion Method Applied to Restricted Diffusion. Simulation of Echo Attenuation from Molecules in Spheres and between Planes. J Magn Reson, Series A, 1993. 104(1): p. 17–25.

62. Ianus, A., et al., Gaussian phase distribution approximations for oscillating gradient spin echo diffusion MRI. J Magn Reson, 2012. 227: p. 25–34.

63. Harkins, K.D., et al., Changes in intracellular water diffusion and energetic metabolism in response to ischemia in perfused C6 rat glioma cells. Magnetic Resonance in Medicine, 2011. 66(3): p. 859–867.

64. Li, H., et al., Impact of transcytolemmal water exchange on estimates of tissue microstructural properties derived from diffusion MRI. Magn Reson Med, 2017. 77(6): p. 2239–2249.

65. Chuhutin, A., B. Hansen, and S.N. Jespersen, Precision and accuracy of diffusion kurtosis estimation and the influence of b-value selection. NMR Biomed, 2017. 30(11): p. e3777.

66. Hui, E.S., et al., Towards better MR characterization of neural tissues using directional diffusion kurtosis analysis. Neuroimage, 2008. 42(1): p. 122–34.

67. Altman, D.G. and J.M. Bland, Measurement in Medicine: The Analysis of Method Comparison Studies.Journal of the Royal Statistical Society: Series D (The Statistician), 1983. 32(3): p. 307–317.

68. Wang, Q., et al., The Allen Mouse Brain Common Coordinate Framework: A 3D Reference Atlas. Cell, 2020. 181(4): p. 936–953.e20.

69. Avants, B.B., et al., The Insight ToolKit image registration framework. Frontiers in neuroinformatics, 2014. 8: p. 44–44.

70. Semple, B.D., et al., Brain development in rodents and humans: Identifying benchmarks of maturation and vulnerability to injury across species. Progress in neurobiology, 2013. 106-107: p. 1–16.

71. Mitchell, S.J., et al., Animal Models of Aging Research: Implications for Human Aging and Age-Related Diseases. Annual Review of Animal Biosciences, 2015. 3(1): p. 283–303.

72. Blumenfeld-Katzir, T., et al., Diffusion MRI of Structural Brain Plasticity Induced by a Learning and Memory Task. PLOS ONE, 2011. 6(6): p. e20678.

73. Dawson, T.M. and T.E. Golde, Animal models of neurodegenerative diseases. 2018. 21(10): p. 1370–1379.

74. Fluri, F., M.K. Schuhmann, and C. Kleinschnitz, Animal models of ischemic stroke and their application in clinical research. Drug design, development and therapy, 2015. 9: p. 3445–3454.

75. Xiong, Y., A. Mahmood, and M. Chopp, Animal models of traumatic brain injury. Nature Reviews Neuroscience, 2013. 14(2): p. 128–142.

76. Zitvogel, L., et al., Mouse models in oncoimmunology. Nature Reviews Cancer, 2016. 16(12): p. 759–773.

77. Praet, J., et al., Diffusion kurtosis imaging allows the early detection and longitudinal follow-up of amyloid-β-induced pathology. Alzheimer’s Research & Therapy, 2018. 10(1): p. 1.

78. Guglielmetti, C., et al., Diffusion kurtosis imaging probes cortical alterations and white matter pathology following cuprizone induced demyelination and spontaneous remyelination. Neuroimage, 2016. 125: p. 363–377.

79. Wirths, O. and T.A. Bayer, Neuron Loss in Transgenic Mouse Models of Alzheimer’s Disease.International Journal of Alzheimer&#x2019;s Disease, 2010. 2010: p. 723782.

80. Blesa, J. and S. Przedborski, Parkinson’s disease: animal models and dopaminergic cell vulnerability.Frontiers in Neuroanatomy, 2014. 8(155).

81. Paxinos, G. and K. Franklin, The Mouse Brain in Stereotaxic Coordinates. 5th ed. 2019: Academic Press.

82. Sampaio-Baptista, C. and H. Johansen-Berg, White Matter Plasticity in the Adult Brain. Neuron, 2017. 96(6): p. 1239–1251.

83. Ianus, A., et al., Accurate estimation of microscopic diffusion anisotropy and its time dependence in the mouse brain. NeuroImage, 2018. 183: p. 934–949.

84. Zhang, H., et al., NODDI: practical in vivo neurite orientation dispersion and density imaging of the human brain. NeuroImage, 2012. 61(4): p. 1000–16.

85. Palombo, M., D.C. Alexander, and H. Zhang. Large-scale analysis of brain cell morphometry informs microstructure modelling of gray matter. in Proc. Intl. Soc. Mag. Reson. Med. 29 2021.

86. Afzali, M., et al., Computing the orientational-average of diffusion-weighted MRI signals: a comparison of different techniques. Scientific Reports, 2021. 11(1): p. 14345.

87. Henriques, R.N., et al., Toward more robust and reproducible diffusion kurtosis imaging. Magnetic Resonance in Medicine, 2021. 86(3): p. 1600–1613.

88. Wu, D., et al., Oscillating gradient diffusion kurtosis imaging of normal and injured mouse brains.NMR in biomedicine, 2018. 31(6): p. e3917–e3917.

89. Henriques, R.N., S.N. Jespersen, and N. Shemesh, Correlation tensor magnetic resonance imaging.NeuroImage, 2020. 211: p. 116605.

90. Henriques, R.N., et al. Measuring the diffusional intra and inter-kurtosis tensors using double diffusion enconding. in Proceedings of the joint meeting of the ISMRM and ESMRB. 2020.

91. Novello, L., et al. Human brain in vivo correlation tensor MRI on a clinical 3T system. in Annual Meeting of the ISMRM. 2021.

92. Yang, D.M., et al., Intracellular water preexchange lifetime in neurons and astrocytes. Magnetic Resonance in Medicine, 2018. 79(3): p. 1616–1627.

93. Lee, H.-H., et al., In vivo observation and biophysical interpretation of time-dependent diffusion in human cortical gray matter. NeuroImage, 2020. 222: p. 117054.

