## Supplementary Information for "Soma and Neurite Density MRI (SANDI) of the in-vivo mouse brain"

*Quality assessment of the data*

Figure S1 presents the maps of the directionally averaged raw signal for b = 1 ms/µm$^2$ and for b = 12.5 ms/µm$^2$ (the highest b-value used in this study) across 32 slices for one representative animal. The first and last slices were excluded due to imaging artifacts. The data is of very high quality and there is a clear WM / GM contrast in the very high diffusion weighted maps. There is also a clear radient in image intensity from the superior to inferior part as data was acquired with a surface coil.

Figure S2 presents the SNR maps of the raw data for all six mice imaged in this study for the two slices illustrated in Figure 3. The SNR values are similar for other slices as well. The SNR was computed as $SNR = \frac{mean(S(b=0))}{std(S(b=0))}$, where *S(b=0)* are the non-diffusion weighted images.

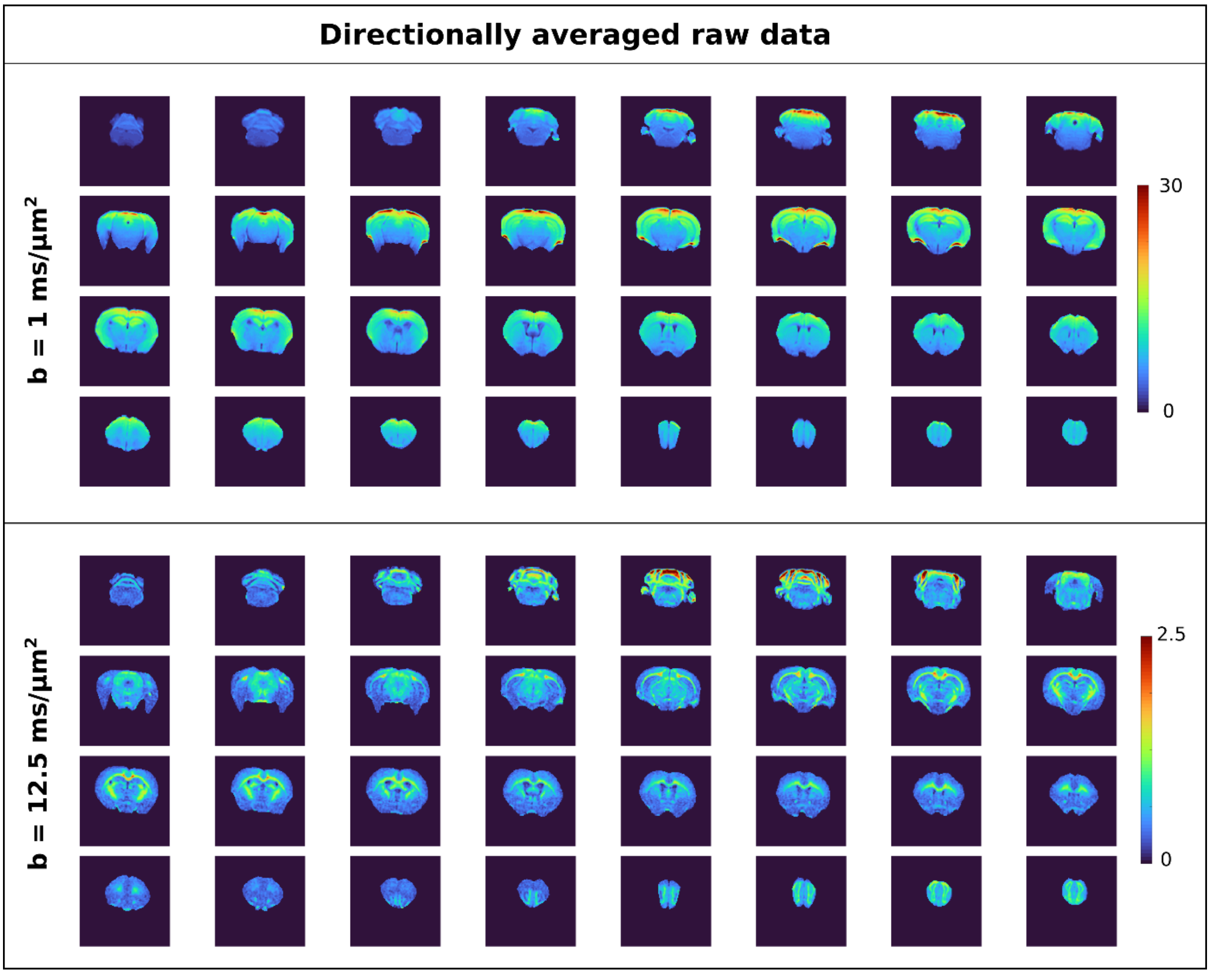


*Figure S 1 Maps of directionally averaged raw data for 32 slices in one representative animal, for b = 1 and 12.5 ms/µm2, which is the highest b-value in the study. The images show a good WM / GM contrast even at the highest b-value.*

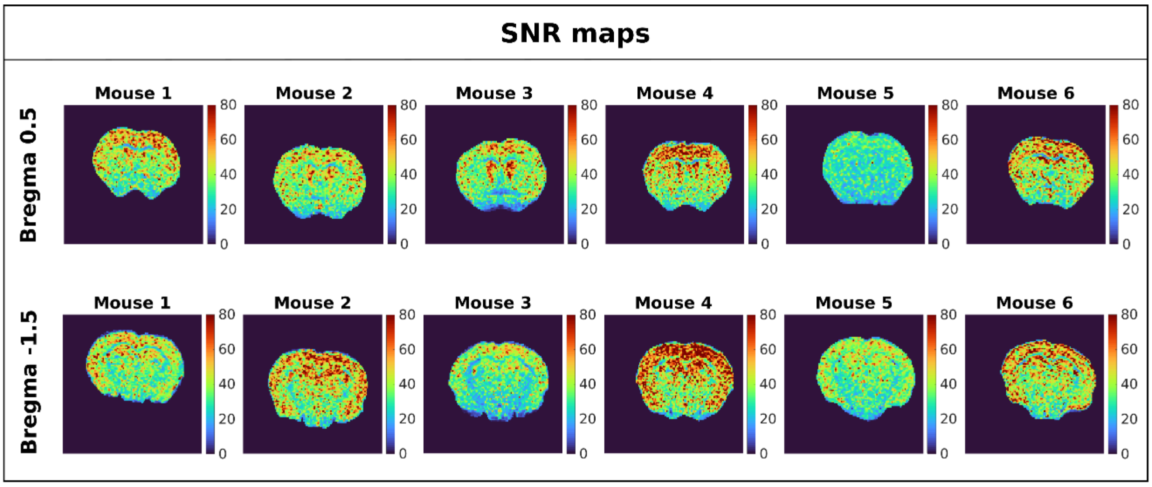


*Figure S 2  SNR maps at different slice positions for all six animals included in this study. There is a slight variation in SNR across animals, nevertheless in all cases SNR > ~30. SNR values are slightly lower in WM, which has lower T2 values and hence lower signal given the relatively long TE of 36.8 ms, as well as in the inferior part of the brain as the images were acquired with a surface coil.*

### *Sandi parameters for magnitude and real data*

This analysis of SANDI parameters estimated from real and magnitude data is complementary to section 3.2. Figure S3 presents the Bland-Altman plots showing the difference between estimates based on magnitude and real data versus the average values. $f_{stick}$ is the parameter which shows the largest positive bias when estimated from magnitude data.

Figure S4 presents the maps of $f_{sphere}$, $f_{stick}$ and $R_{sphere}$ for 32 slices in one representative animal. We see that the contrast between WM and GM is consistent across slices.

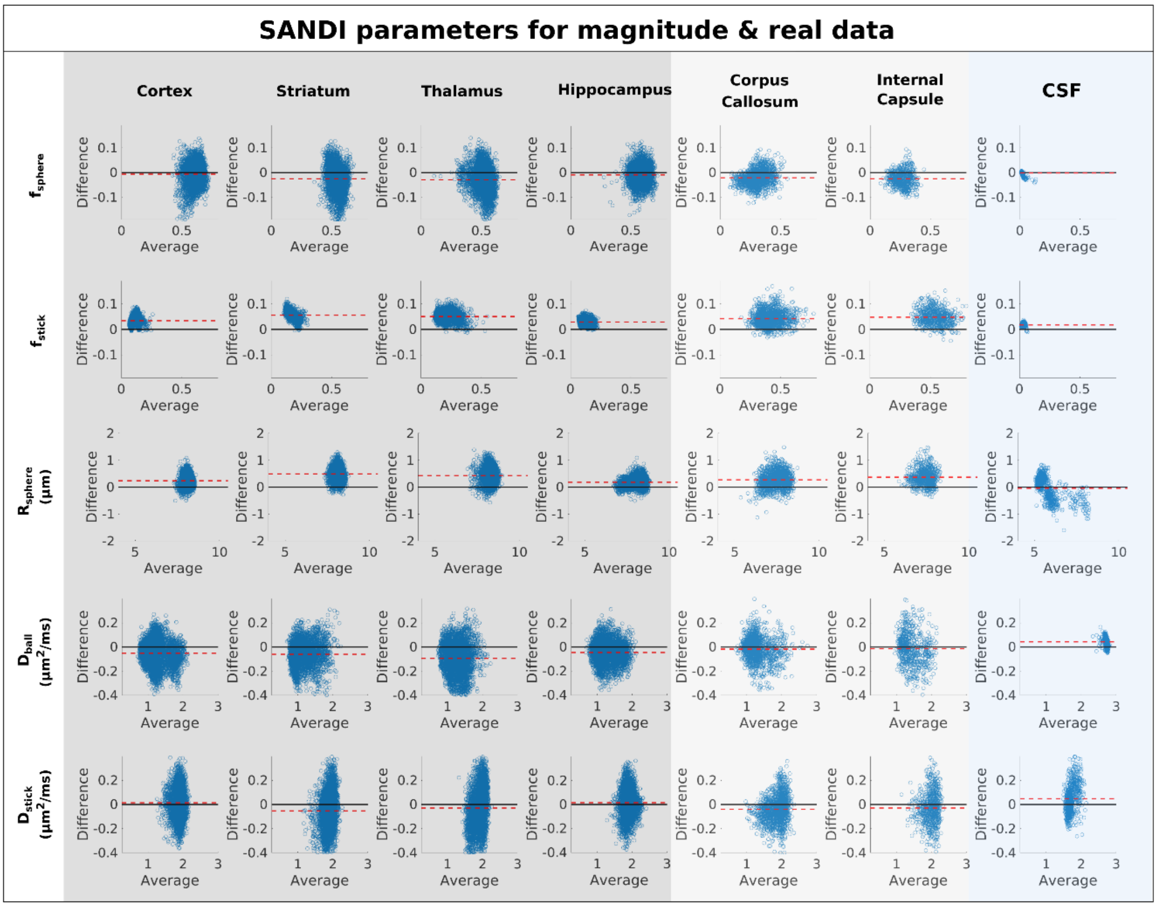


*Figure S 3 Bland-Altman plots showing the difference in parameter values estimated from magnitude and real data plotted as a function of their average value. The dotted red line represents the mean difference, while the black line shows the zero value.*

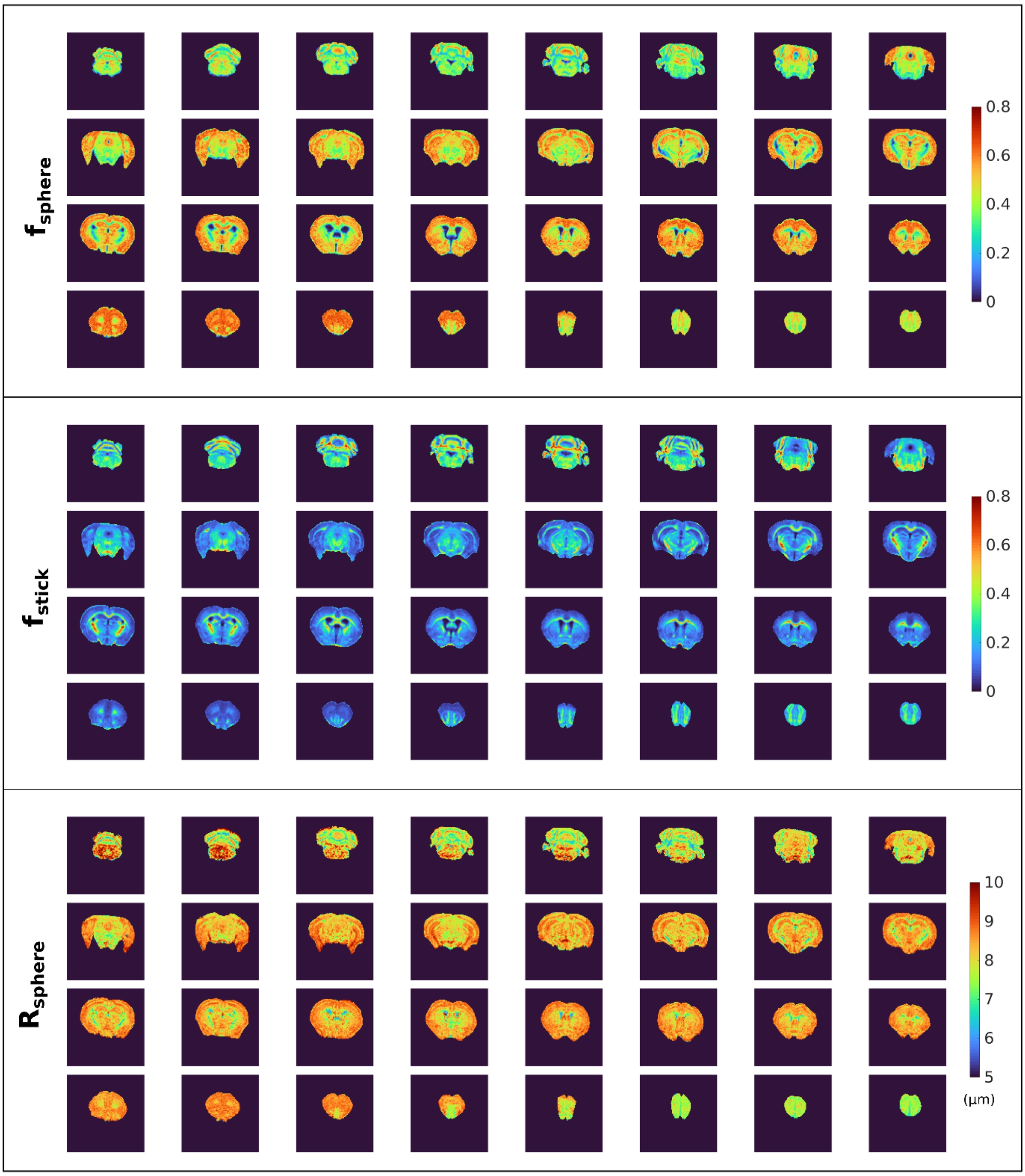


*Figure S 4 Brain wide maps of* $f_{sphere}$, $f_{stick}$ *and* $R_{sphere}$ *for 32 slices in one representative animal*

*dMRI parameter values in GM, WM and CSF ROIs*

Table S1 presents the mean and standard deviation across mice for the ROI averaged SANDI and DKI parameter values corresponding to the histograms in Figure 6.

| | Cortex | Striatum | Thalamus | Hippocampus | Corpus Callosum | Internal Capsule | CSF |
|---|---|---|---|---|---|---|---|
| $f_{sphere}$ | 0.62±0.02 | 0.55±0.03 | 0.52±0.02 | 0.59±0.02 | 0.32±0.01 | 0.27±0.02 | 0.02±0.005 |
| $f_{nstick}$ | 0.09±0.01 | 0.15±0.01 | 0.18±0.02 | 0.10±0.004 | 0.40±0.03 | 0.52±0.02 | 0.01±0.002 |
| $D_{ball}$ $(\mu m^2/ms)$ | 1.26±0.18 | 1.21±0.11 | 1.32±0.12 | 1.33±0.19 | 1.32±0.08 | 1.43±0.16 | 2.68±0.01 |
| $R_{sphere}$ $(\mu m)$ | 8.03±0.06 | 7.88±0.15 | 8.06±0.09 | 8.04±0.06 | 7.29±0.14 | 7.19±0.18 | 5.88±0.25 |
| $D_{stick}$ $(\mu m^2/ms)$ | 1.88±0.05 | 1.90±0.06 | 1.92±0.05 | 1.87±0.06 | 1.86±0.06 | 1.98±0.06 | 1.70±0.02 |
| MD $(\mu m^2/ms)$ | 0.69±0.04 | 0.68±0.03 | 0.72±0.03 | 0.70±0.04 | 0.70±0.03 | 0.71±0.03 | 3.3±0.15 |
| FA | 0.15±0.02 | 0.20±0.04 | 0.27±0.04 | 0.16±0.03 | 0.60±0.04 | 0.59±0.07 | 0.13±0.04 |
| MK | 0.52±0.09 | 0.56±0.06 | 0.57±0.09 | 0.50±0.12 | 0.63±0.11 | 1.01±0.14 | 0.26±0.04 |

*Table S 1 Mean and standard deviation across animals for SANDI and DKI parameters in different ROIs.*

*DKI parameter maps*

Figure S5 illustrates DKI parameter maps, for the same representative animal and slices shown in Figure 4 for the SANDI parameters.

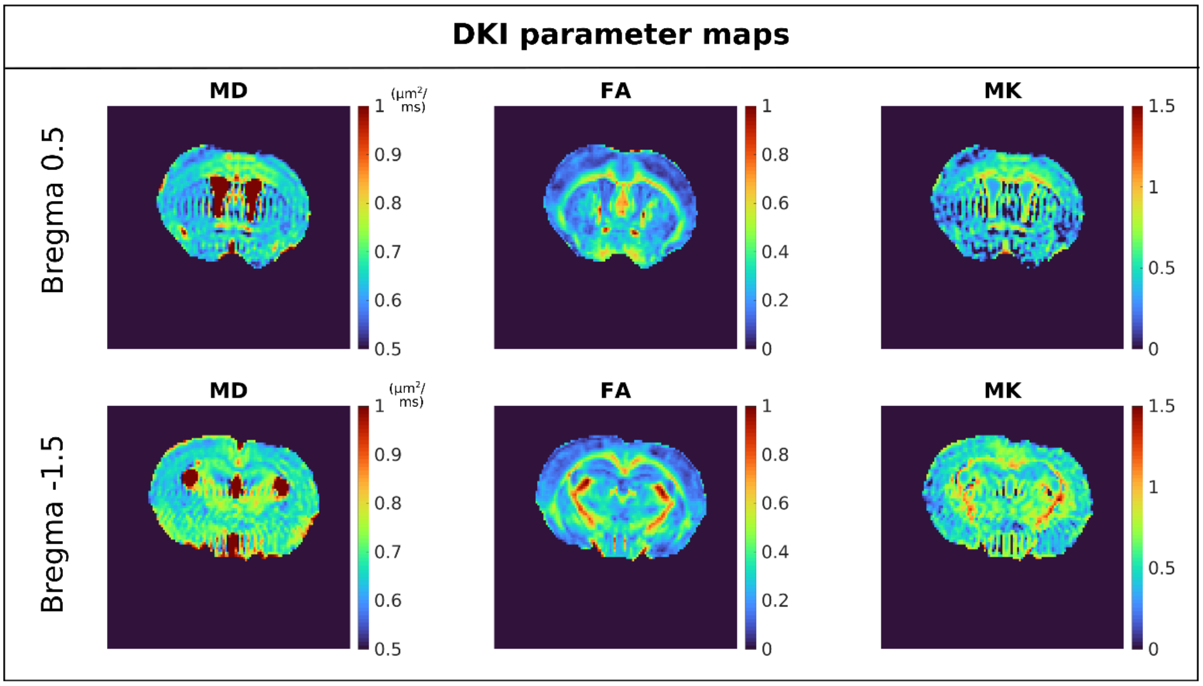


*Figure S 5 DKI parameter maps in one representative mouse*

Variation of $f_{sphere}$ across mice

Figure S6 presents maps of $f_{sphere}$ for all the mice imaged in this study, for the slices used to study the correlation with the image intensity of the Allen mouse brain atlas.

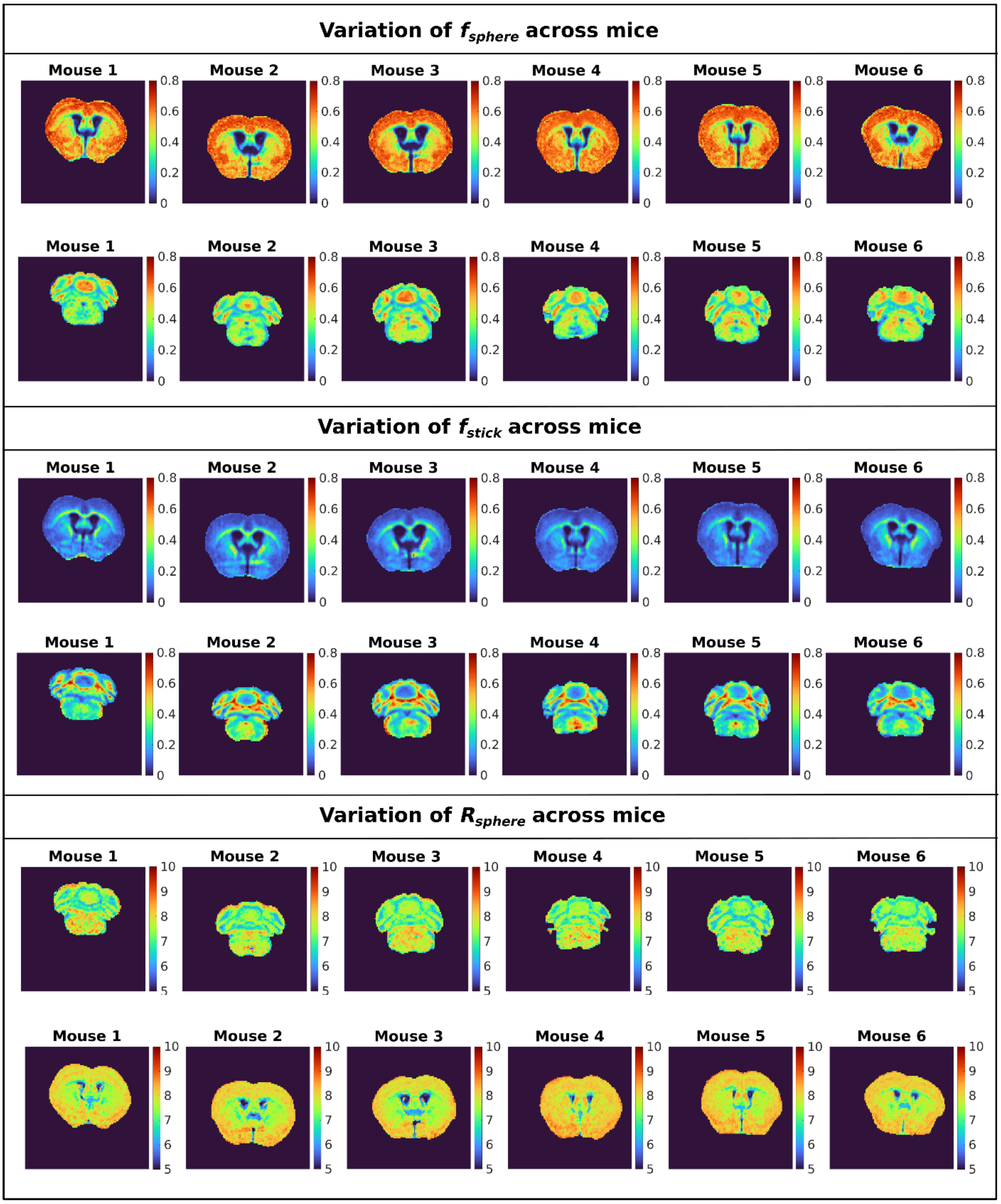


*Figure S 6 Maps of $f_{sphere}$ for two slices in cerebrum and cerebellum for all mice included in the study, prior to registration.*

Variation of $R_{sphere}$ gray matter ROIs

Figure S7 shows maps of the $R_{sphere}$ parmeter in three slices of a representative mouse after a slight smoothing with a 2D Gaussian kernel with standard deviation of 0.5 to better illustrate variations in $R_{sphere}$ across the brain.  Larger $R_{sphere}$ values are estimated in areas such as the pyramidal layer of the hippocampus (black arrows) and the piriform cortex (white arrows).

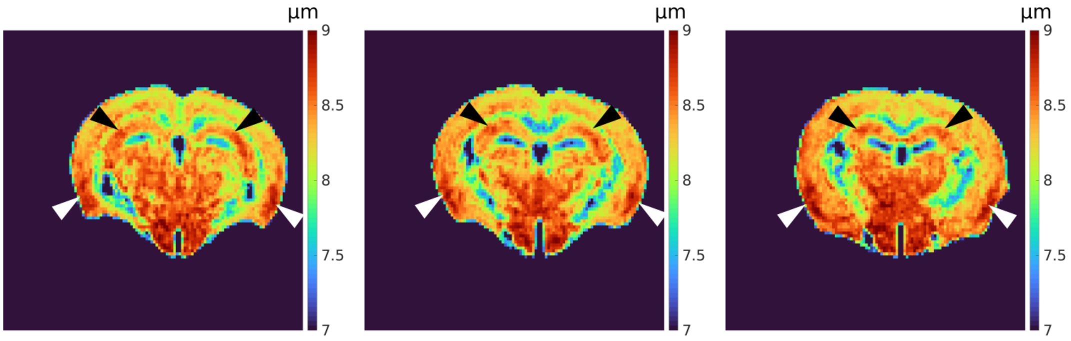


*Figure S 7 Maps of* $R_{sphere}$ across three slices in a representative mouse after a slight smoothing with a 2D Gaussian filter with standard deviation 0.5. Higher $R_{sphere}$ values are estimated in the pyramidal layer of the hippocampus (black arrows), the piriform cortex (white arrows), as well as in the thalamus and hypothalamus.
